# Large-scale analysis of *Drosophila* core promoter function using synthetic promoters

**DOI:** 10.1101/2020.10.15.339325

**Authors:** Zhan Qi, Christophe Jung, Peter Bandilla, Claudia Ludwig, Mark Heron, Anja Sophie Kiesel, Julia Philippou-Massier, Miroslav Nikolov, Alessio Renna, Max Schnepf, Ulrich Unnerstall, Johannes Soeding, Ulrike Gaul

## Abstract

The core promoter, the region immediately surrounding the transcription start site, plays a central role in setting metazoan gene expression levels, but how exactly it ‘computes’ expression remains poorly understood. To dissect core promoter function, we carried out a comprehensive structure-function analysis to measure synthetic promoters’ activities, with and without an external stimulus (hormonal activation). By using robotics and a dual-luciferase reporter assay, we tested ∼3000 mutational variants representing 19 different *Drosophila melanogaster* promoter architectures. We explored the impact of different types of mutations, including knockout of individual sequence motifs and motif combinations, variations of motif strength, positioning, and flanking sequences. We observe strong effects of the mutations on activity, and a linear combination of the individual motif features can largely account for the combinatorial effects on core promoter activity. Our findings shed new light on the quantitative assessment of gene expression, a fundamental process in all metazoans.

## INTRODUCTION

Appropriate gene expression with the correct timing is crucial for the development and diversity of all organisms. The control of gene expression occurs primarily at the process of transcription (Levine and Tjian, 2003), and the core promoter makes an essential contribution for setting the gene expression level (Lubliner et al., 2015).

The RNA polymerase II (Pol II) core promoter is the minimal DNA sequence that is recognized by the basal transcription machinery (Juven-Gershon et al., 2008; Smale and Kadonaga, 2003; Thomas and Chiang, 2006). It comprises the transcription start site (TSS) and approximately 150 bp of the flanking sequence. The accurate transcription initiation and basal expression level of a gene are primarily determined by differential recruitment of the transcription machinery (consisting in Pol II and general transcription factors (GTFs)) to its core promoter region (Juven-Gershon et al., 2008; Smale and Kadonaga, 2003; Thomas and Chiang, 2006). Genome-wide studies have revealed various properties of native core promoters. In particular, sequence motifs that are over-represented around TSSs mostly mark the potential binding sites of GTFs or other transcription factors (TFs) (Burke and Kadonaga, 1997; FitzGerald et al., 2006; Ohler, 2006; Parry et al., 2010b). Genetic variations occurring at the motif sites alter both promoter strength and TSS position significantly (Schor et al., 2017). Although the genomic analysis of native sequences suggests certain causal relationships, the variations in genomic sequences have been very challenging to predict (Seizl et al., 2011). This makes it difficult to uncover the sequence attributes responsible for activity changes. Hence, it remains impossible to ascertain the influence of specific features except by directly altering them and measuring the effect on expression levels.

Facilitated by DNA synthesis technology and next-generation sequencing, high-throughput approaches such as massively parallel reporter assays (MPRAs) have been developed to test how the DNA sequence affect gene expression on a large scale (Arnold et al., 2013; Melnikov et al., 2012; Patwardhan et al., 2009; Sharon et al., 2012). The dynamic range of the RNA-seq based MPRAs is usually small (around two orders of magnitude) and they suffer from low accuracy for weak regulatory elements that have a low read coverage. These limitations would severely influence a core promoter analysis, because it is known to drive basal and modest expression. A second kind of MPRA method quantifies the protein fluorescence as the readout of reporter gene expression but can only obtain discrete expression measurements because of their “bin” sorting design (which cannot sense subtle effects), and of the intrinsically narrow dynamical range of the fluorescence readout (Lubliner et al., 2015). Finally, most of the studies focus on enhancers, especially on TF binding sites. Only few MPRAs were designed for *in vivo* promoter analysis, such as extensive studies on fully designed yeast proximal promoter regions (Sharon et al., 2012) and yeast core promoter sequences (Lubliner et al., 2015), analysis of autonomous promoter activity of random genome fragments in human (Van Arensbergen et al., 2017), and in *Drosophila melanogaster (D. mel.)* (Arnold et al., 2017). In the latter study, the measurements of the STAP-seq method were unfortunately not sensitive enough to study the basal activity of the putative promoters in *D. mel.,* but only their enhancer responsiveness. Thus, despite the pivotal role of core promoters in transcription initiation, it remains poorly understood how the components and sequence features of the core promoter determine expression levels.

This study aims to dissect the core promoter comprehensively and to elucidate the sequence determinants of promoters in *D. mel.* S2 cells. We test promoter activity using a dual luciferase assay, which is highly sensitive with a linear and broad dynamical range. We have integrated the entire experimental pipeline using automated robotic systems, including cloning and luciferase gene expression readout (Figure 1 and S1). This allowed us to accurately measure promoter activity at large scale and with high reproducibility. By extensively testing mutagenized core promoter sequences for 19 genes, we corroborate the functional specificity of sequence motifs. We demonstrate that their strength, as measured by the position weight matrix (PWM) score, and their precise positioning are essential features determining core promoter activity. Additionally, we comprehensively mutagenized core promoter motifs using single base pair mutations to produce expression-based position probability matrices (PPMs) and activity logos for them. Combinatorial motif mutations that alter both the strength and the positioning of all motifs often result in strong effects on activity, which are compared with the effects of individual motif mutations: we find that a linear combination of these individual motif features can largely account for the joint effects on core promoter activity. In addition, we investigate the influence of surrounding promoter regions on promoter activity, especially for the ecdysone response element (EcRE). The ecdysone responsiveness depends on the core promoter architecture. Ecdysone can induce both developmental and constitutive core promoters but the induction is stronger with the developmental ones. We also find a negative correlation between the ecdysone inducibility and the basal expression level; this correlation is more significant for constitutive promoters. Finally, by testing sequences impacting −1 and +1 nucleosomes, we found that their influence on constitutive core promoter activity is relatively mild, the effect being stronger for nucleosome positioning sequences downstream of the TSS.

**Figure 1.**
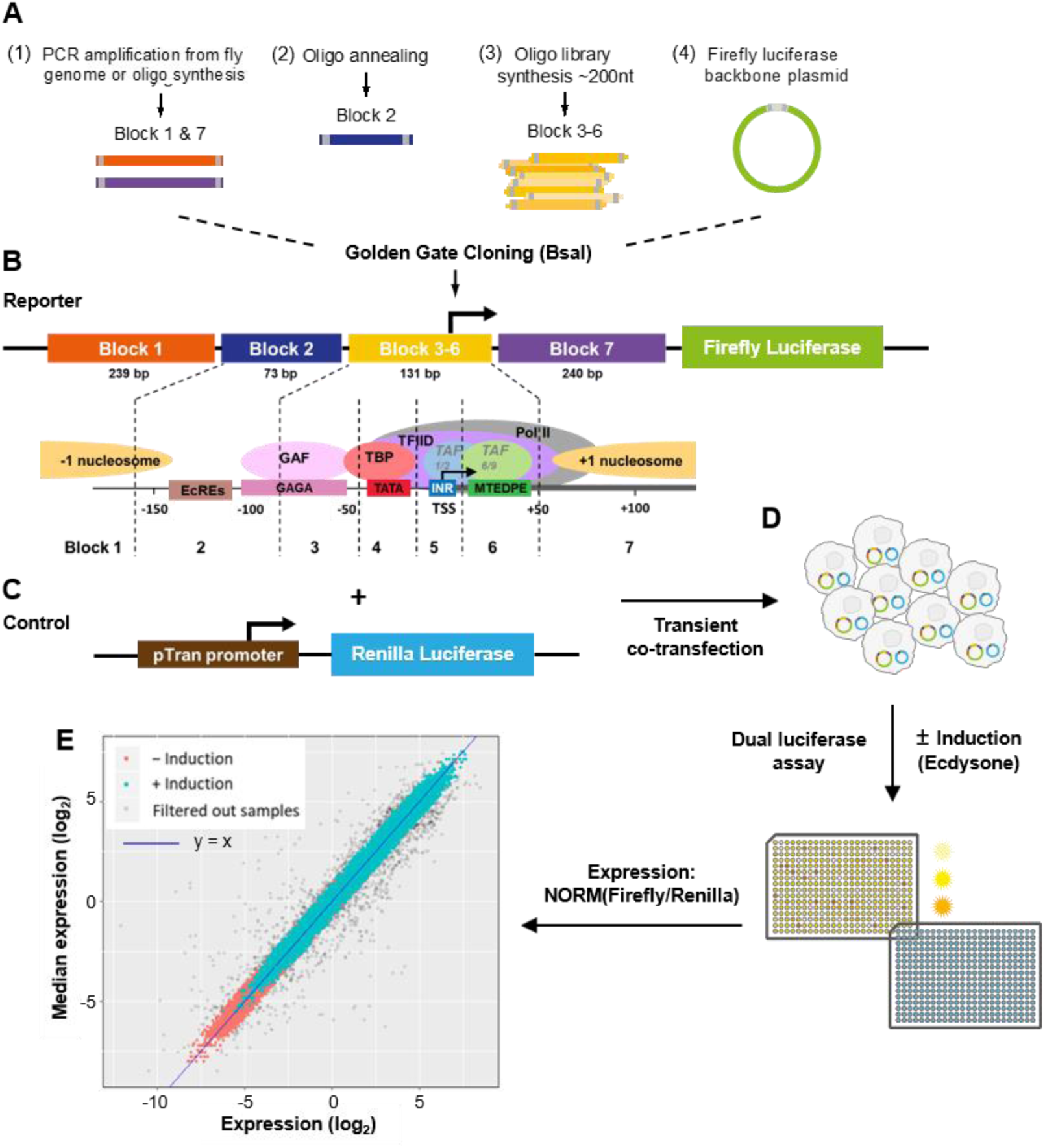
Experimental workflow and assay reproducibility. (A) The promoter region was divided into 7 building blocks: *block 1* with 239 bp of a potential −1 nucleosomal sequence; *block 2* with 73 bp sequence representing the ecdysone receptor binding region; *block 3-6* with 131 bp sequence representing the native and perturbative core promoter regions from different architectures; *block 7* with 240 bp of a potential +1 nucleosomal sequence. (B-D) Simplified dual luciferase assay experimental workflow. To measure promoter activity quantitatively on a large scale with high reproducibility we integrated the golden gate cloning strategy (BsaI cloning) with a high-throughput experimental pipeline using automated robot systems for colony picking, reporter plasmids isolation, transient co-transfection and dual luciferase assay (**MATERIAL AND METHODS** for details). (E) Reproducibility of normalized expression levels for all tested promoters. The expression levels covered a wide dynamic range of more than four orders of magnitude. Expressions with and without ecdysone induction are labeled cyan and red, respectively. About 5% of the raw data were filtered out as outliers (gray dots; see **MATERIAL AND METHODS** for details). Blue line: y = x.

## RESULTS

### Design and selection of synthetic promoter sequences

Our design intends to test three different promoter features separately: core promoter sequence features (especially motifs), transcriptional response to external stimulus, and influence of genomic ±1 nucleosomal sequences. The tested promoter sequences were inserted into synthetic constructs made of combined building sequence blocks, which comprise different functional regions (Figure 1A and 1B): (1) a motif-rich core promoter region of 130 bp around the TSS with native and perturbed sequences from different core promoter architectures (referred to as *block 3-6);* (2) a stimulus-response element for binding of the ecdysone receptors to recruit the steroid hormone ecdysone for transcriptional activation *(block* 2); and (3) genomic −1 and +1 nucleosome positioning sequences to mimic the endogenous ±1 nucleosomal context *(block 1* and 7, respectively).

We defined the genes to be tested based on experimentally genome-wide derived features, including expression strengths and variation during developmental stages (Graveley et al., 2011), Pol II stalling (Hendrix et al., 2008; Zeitlinger et al., 2007), TSSs mapping from CAGE data (Hoskins et al., 2011; Ni et al., 2010), and motif composition. We applied *XXmotif* algorithm (Luehr et al., 2012) for a genome-wide *de novo* motif search in annotated core promoter regions and were able to identify widely known motifs as well as some novel motif candidates with optimized PWMs based on enrichment, localization and conservation (**Table S1**). By correlating all identified motifs to the gene sets (**Figure S2A - S2B**), four architectures of core promoter motifs could be defined (Figure 2 and S2C): two architectures were attributed to developmental promoters (7 promoters selected in this study), the two other architectures to constitutive ones (9 selected). We also found an additional class of promoters containing no known motifs (3 selected). For the developmental and constitutive promoters (Figure 2 and S2C), all identified motifs are known motifs including INR, MTE/DPE (an overlapping version of the two previously identified motifs MTE and DPE, hereafter referred to as MTEDPE), GAGA, GAGArev, INR2 (widely known as motif 1 or Ohler1), DRE, Ohler7, E-Box1, Ohler6, TATA-Box, R-INR (widely known as TCT motif, here named as ribosomal initiator based on its co-localization with TSSs of ribosomal protein genes), E-Box2; we named the new motifs CGpal, INR2rev, TTGTT, TTGTTrev, AAG3, ATGAA and RDPE (ribosomal downstream promoter element). In particular, another new motif named CA-INR was often found co-occurring with TATA-Box, which is a highly conserved derivative of the classical INR motif. CA-INR also has a strong positional preference around TSS and its most representative sequence is GGCATCAGTC with the TSS mostly mapped at its 4^th^ position.

**Figure 2.**
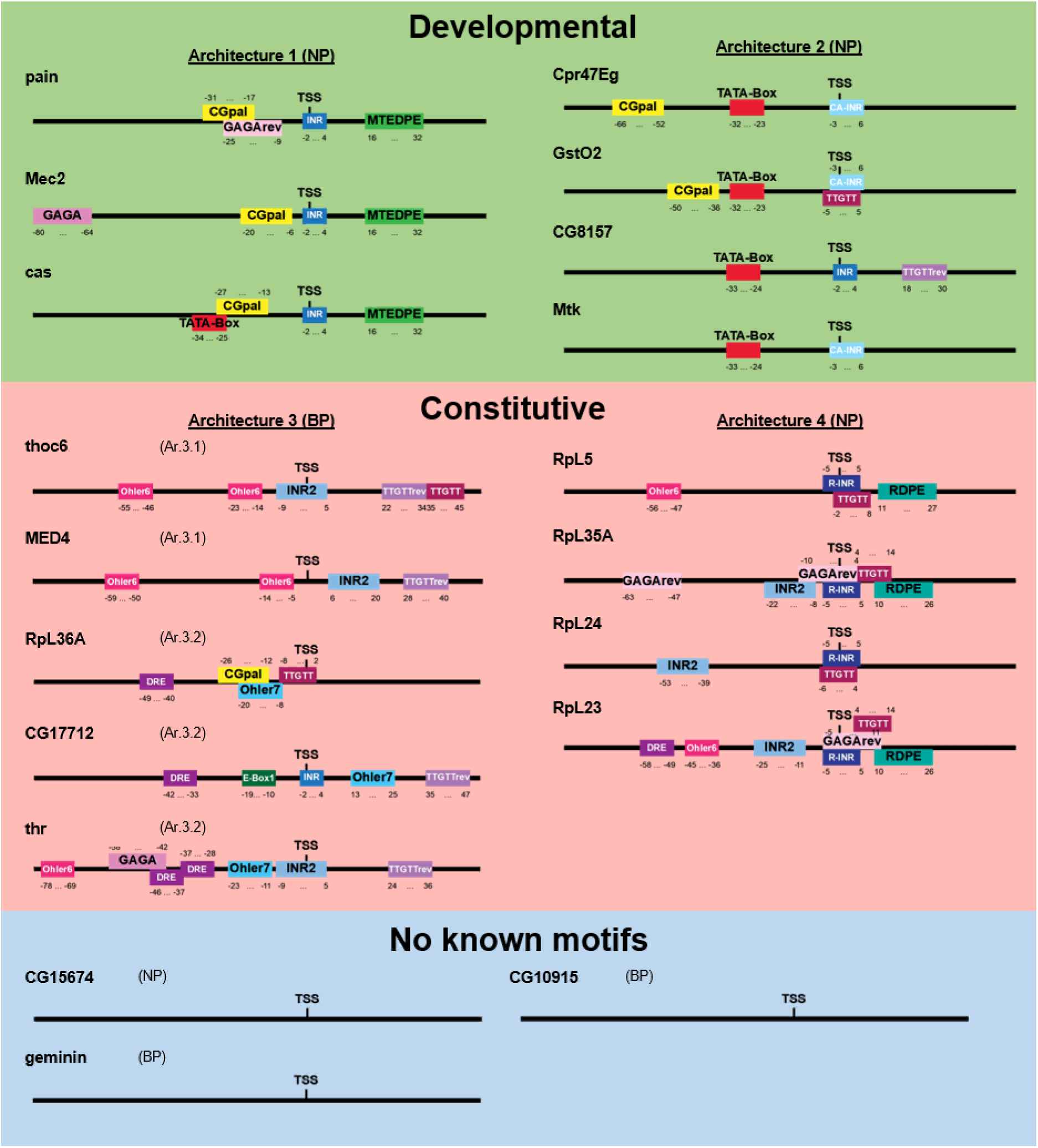
The wild-type core promoters and their motif composition. (A) From each of the four core promoter architectures Ar.1, Ar.2, Ar.3 (Ar.3.1, Ar.3.2), Ar.4 and one additional architecture with no known motif (termed motif-less promoters), 2-4 native sequences were chosen (position −80 to +50 relative to TSS; TSS itself at position 0). In total 19 wild-type core promoters with annotated motif positions are shown here. NP, narrow peak; BP, broad peak. Their sequences are listed in **Table S3**. Developmental and constitutive promoters are highlighted in green and red, respectively. Motif-less promoters in blue.

To systematically examine the sequence motifs of the motif-rich core promoter *(block 3-6)* we devised various mutations of wild-type promoters (Figure 3), including individual or pairwise knockout (complete replacement with non-functional sequences) of motifs, knockout of all motifs, replacing the original motif with its *XXmotif*-derived highest frequent genomic sequence (hereafter referred to as consensus), point mutations of motifs, shift of motif positions and substitution with functionally or positionally equivalent motifs from other architectures. In addition to widely known motifs like INR and TATA-Box, we also tested four of the new motif candidates discovered by *XXmotif* (CGpal, TTGTT, TTGTTrev, and RDPE; **Tables S1** and **S2**). We compared the activities measured from synthetic promoters containing mutated motifs with the corresponding wild-type strengths. The results obtained with the point mutations allowed an analysis of motif specificity. Recent studies on TF binding suggest that the sequence motifs alone cannot fully explain the activity variation (Schone et al., 2018; Yella et al., 2018). Therefore, we also tested in our experiments the context sequences surrounding the motifs. Finally, combinatorial mutations altering both strength and positioning of all motifs within core promoter architectures as well as block-wise swaps between architectures were performed for more in-depth analysis which enabled quantitative modeling of promoter activity based on individual sequence features.

**Figure 3.**
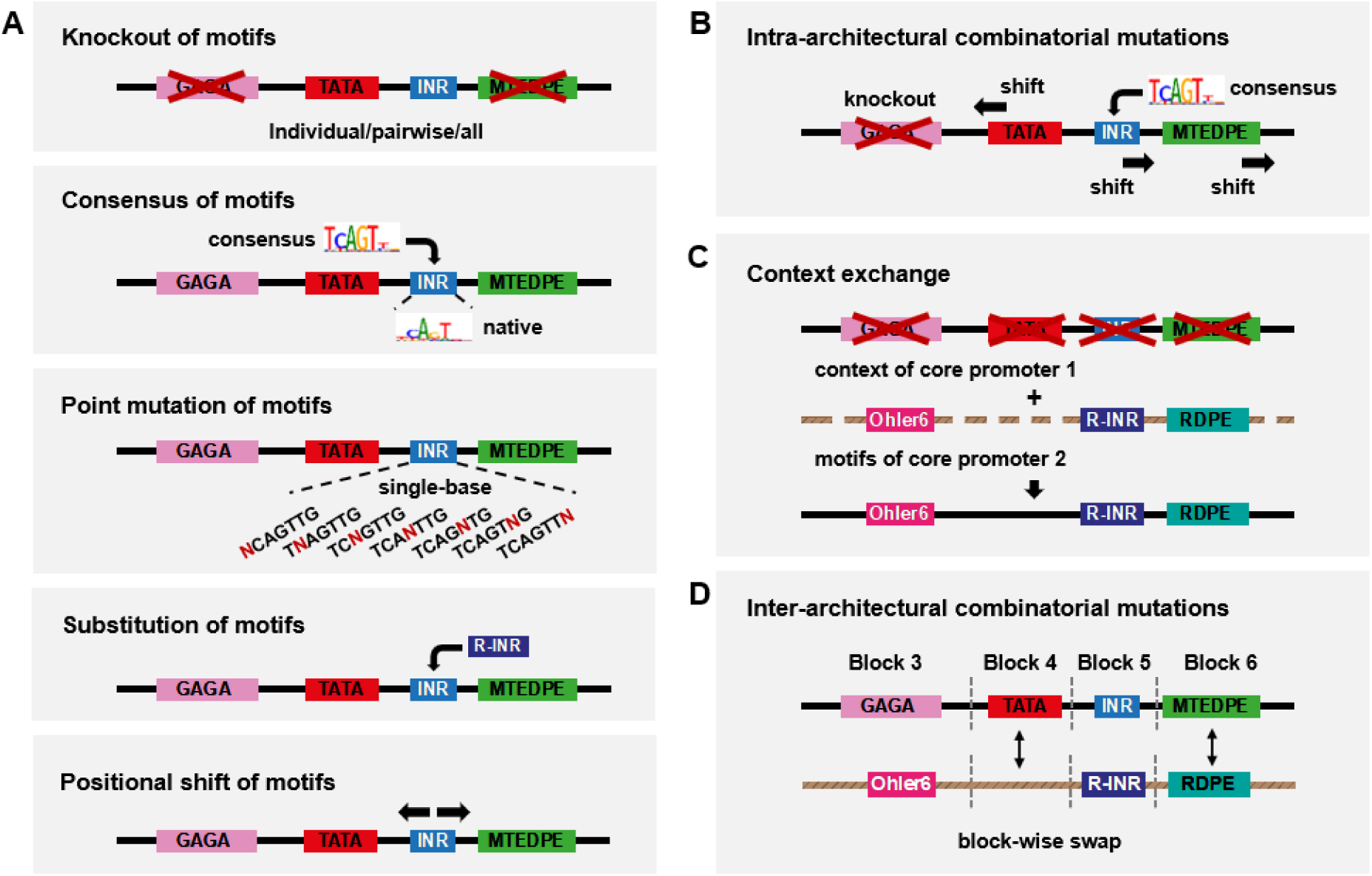
Combinatorial mutations designed for the motif-rich core promoter region. (A) Motif-wise combinatorial mutations within the core promoter. Motif strength and motif position are changed individually: knockout of motifs (individual or pairwise knockout of motifs, and knockout of all motifs); replacing the original motif with its computationally *(XXmotif)* derived sequences with different PWM scores (consensus with the highest score), or insertion of the consensus into the motif-less promoter sequences; point mutation of motifs; substitution with functionally or positionally equivalent motifs from other architectures; shift of motif positions. The Mec2 motif composition is shown here as an example. (B) Intra-architectural mutations: both change of motif strength and motif position within the same construct. (C) Context exchange between different core promoters. (D) Inter-architectural mutations: block-wise combinatorial mutations between different core promoters.

### Core promoter activity measurements for thousands of designed sequences

We applied our method to produce and measure both basal and induced expressions of synthesized oligonucleotides representing wild-type (**Table S3**) and mutated core promoters. We designed in total 3826 synthetic promoter sequences (**Table S4**), and were able to recover and test experimentally ∼3000 of these sequences (**MATERIAL AND METHODS**). The *block 3-6* sequences were assembled with one inducible *block 2* and different combinations of *block 1* and *block 7* nucleosomal sequences, constructing the entire library of synthetic promoters to be tested in our experiments (Figure 1). For most of the constructs (> 88%), we measured with and without ecdysone stimulation at least three replicates each. The expression levels range over more than four orders of magnitude (Figure 1, lower left panel) and have a very high reproducibility among replicates, with a mean coefficient of variance (CV) of 21% (Figure S1D; median standard deviation of 29%).

All *block 1*s and *block 7*s are native *D. mel.* nucleosomal sequences selected to provide a variety of nucleosome occupancies (determined using our own MNase digestion followed by high-throughput sequencing (MNase-Seq) dataset, data not shown). Probing with our assay the pair-wise *block 1.X* and *7.X* (with X an arbitrary index corresponding to the gene selected for their nucleosomal sequences; **Table S5** and **S6**) showed that the paired *block 1.11* and *block 7.11* (hereafter termed as *B1.11* + *B7.11)* gave the highest expression (**Figure S3A**). We checked by MNase-Seq the nucleosome occupancy on the plasmid of the synthetic promoter construct containing this pair (**Figure S3B**, higher panel) and nucleosome patterns were visible on the *B1.11* and *B7.11* sequences; they were similar to what was observed at the genomic locus (**Figure S3B**, lower panel). Therefore, this *B1.11* + *B7.11* combination was selected as the fixed nucleosomal context sequence for highly mutated *block 3-6*s in the subsequent experiments. We systematically tested combinations of different *block 1s* and *7s* (**Figure S3C - S3E** and **Supplemental Text 1**): different potential nucleosomal contexts showed moderate effects on expression levels with a stronger effect for sequences potentially forming +1 nucleosomes. Constitutive core promotes were more sensitive to the influence of nucleosomal sequences downstream of the TSS.

To determine the activity level range of the native core promoters (**Table S4**), we measured the constructs containing all wild-type (i.e. native) promoter sequences with *B1.11* + *B7.11.* The expression levels showed a broad range that spanned over three orders of magnitude (Figure 4A). Two housekeeping core promoters MED4 and CG17712 drove the highest expressions, while the ribosomal class generally showed an intermediate activity. As expected, the core promoters with no known motif showed the lowest activity (in blue in Figure 4A).

(D) Mean expression fold changes compared to wild-type expressions for individual knockout of motifs in different core promoters. Constitutive and developmental promoters are highlighted in red and green, respectively.

(E-F) Effect of pairwise motif knockout (log_2_ scale) in core promoters CG7712 (E) and pain (F), respectively. The heatmaps display the mean expression fold changes compared to wild-type expressions for pairwise knockout of motifs compared to individual knockouts (diagx00D7;onals). Additivity was calculated as the difference between the pairwise effect and the sum of two individual effects, Subadditive (in blue): > 0; superadditive (in yellow): < 0; Additivity values for effects > 3×SD_noise_ shown in the right lower corner of each pairwise effect.

**Figure 4.**
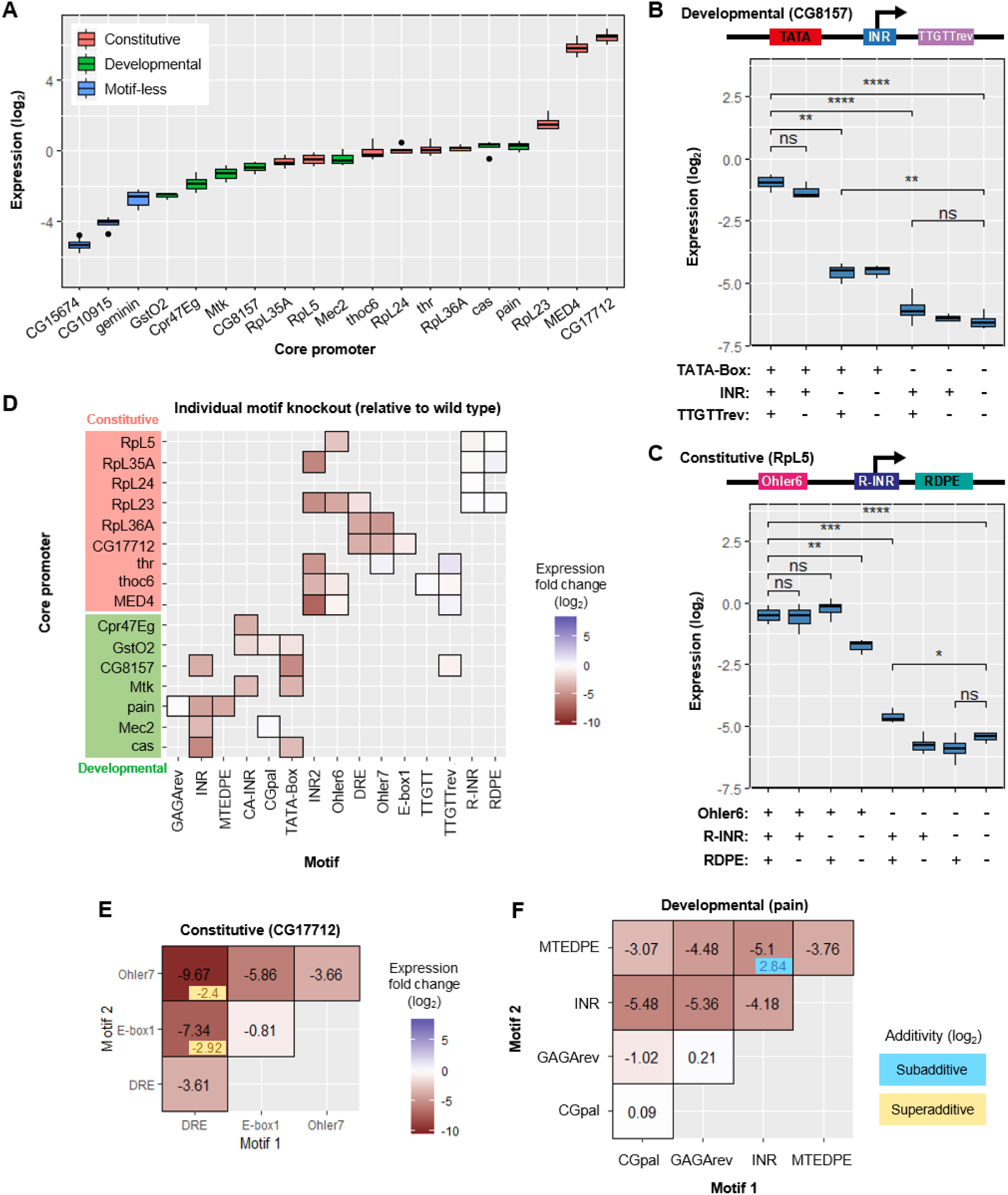
Motif knockout. (A) Normalized expression levels of the native core promoters. Their activities spanned a broad range (over three orders of magnitude; promoter constructs contained *block 1.11* and *block 7.11* combination as nucleosomal sequences). Each color represents a different class of core promoter architecture. (B-C) Comparison of normalized expression levels between wild-type configuration and motif knockouts for two types of core promoters (developmental: CG8157; constitutive: RpL5). Upper panels: schematic depiction of the wild-type motif compositions (TTGTT motif in RpL5 is ignored due to its strong overlap with R-INR). Two-sample t-test: ns, not significant, p > 0.05; *p < 0.05; **p < 0.01; ***p < 0.001; ****p < 0.0001.

### Knockout of motifs generally decreases expression consistently between core promoters

To find out whether motif knockouts significantly affect expression, we compared the expression levels of the wild-type configuration with individual, pairwise, and all-motif knockouts (Figure 1B - 1F and **S4A - S4G**). The disruption of well-known motifs like INR and TATA Box in CG8157 (Figure 4B) or Ohler6 in RpL5 (Figure 4C) reduce activity substantially. The only exception is the initiator for the ribosomal protein genes (R-INR) that showed no significant effect when mutated in RpL5 (Figure 4C) or in any other tested ribosomal core promoter (Figure 4D). A similar absence of effect was observed for the RDPE motif, while a knockout of both motifs did cause a decrease in expression in RpL5 (∼ 2.4-fold reduction; Figure 4C). Disrupting all motifs in both promoters led to much weaker expressions (> 30-fold decrease) (Figure 4B and 4C).

More generally, the knockout of all motifs resulted in a near complete loss of function for each tested core promoter sequences, regardless of its wild-type strength (**Figure S4A**). Most of these all-motif knockout configurations exhibited lower activity than wild-type promoters containing no known motif (**Figure S4A**). Compared to wild-type expression, knocking out individual motifs typically resulted in a reduction (Figure 4D). These effects were consistent across the different promoters. An exception is the knockout of TTGTTrev that slightly increased expression in Thr and MED4 (the blue in the middle right of Figure 4D; > 2-fold increase after disruption of the motif). Hence, this motif functioned as a weak repressor in these promoters. The core of TTGTTrev (AACAA) matches the central part of the binding site of an adult enhancer factor (AEF-1) in *D. mel.,* which is known to be a short-range transcriptional repressor (Brodu et al., 2001; Falb and Maniatis, 1992a, b). Finally, the ribosomal promoter motifs R-INR and RDPE did not lead to a reduction of activity after the disruption in all the four investigated constitutive promoters (top right corner in Figure 4D).

### Pairwise knockouts of some motifs show synergistic (superadditive) effects

To investigate the role of motif interplay on regulating the expression, we compared the results obtained from pairwise knockouts with their individual knockout measurements in different core promoter configurations. Overall, the effect of most pairwise knockouts was additive (in log scale; Figure 4E - 4F and **S4B – S4G**). However, in some cases the expression levels were greater or less than the sum of the individual effects (super-and sub-additive effects, respectively). For instance the motif pairs DRE + Ohler7 and DRE + E-Box1 in promoter CG17712 showed strong synergistic interactions (Figure 4E): the double knockouts yielded respectively a 2^2.4-^ fold and 2^2.9^-fold lower expression than the repression expected from their independent, added effects on log2 expression. DRE is considered the most crucial motif in this housekeeping core promoter architecture as it directs a specific TF DREF binding (Hirose et al., 1993). The strong superadditivity we observed suggests the existence of a compensatory phenomenon for DREF binding involving Ohler7 and/or E-Box1 against potential mutations of the DRE motif. Ohler7 could fully recover the activity when E-Box1 was disrupted, but not *vice versa* (CG17712 in Figure 4E). Nevertheless, core motifs in developmental promoters such as INR and MTEDPE in the pain promoter (Figure 4F), or INR2 and Ohler6 in the RpL23 promoter (Figure S4F) are so crucial for expression activity that a knockout of either resulted in almost the same effect as disrupting them both (subadditivity). For promoters GstO2, thoc6 and MED4, the pairwise effects showed exclusively linear additivity (**Figure S4B -– S4D**).

Taken together, these results demonstrate that the disruption of some motif pairs in a given core promoter leads to synergistic effects. DRE is crucial for housekeeping promoter function, and the other three housekeeping motifs Ohler6, Ohler7 and INR2 also play essential roles in regulating ribosomal gene transcription.

### Most motif consensus sequences drive higher expression

In addition to motif knockout, we tested if computationally derived consensus sequences that are preferred in the genome could increase expression (Figure S4H). Most consensus sequences drove higher promoter activity, especially the consensus of TATA-Box in GstO2 (more than 15-fold stronger expression; seen as dark blue square in Figure S4H). As an exception, replacing the TTGTTrev motifs with their consensus sequence in three promoters led to a signal reduction, again supporting its role as repressor (brown square in **Figure S4H**).

Because replacing most motifs with their consensus sequence increased expression levels, we asked whether these sequences could boost the activity of the motif-less promoters (CG15674, CG10915 and geminin) (Figure 5A). Indeed, some motifs, particularly those containing a CA TSS site like INR, INR2, and Ohler7, were sufficient to significantly induce expression when inserted into these motif-less promoters (Figure 5A and 5B; > 2-fold increase for INR replacement, ∼ 100-fold increase for INR2 and ∼ 5-fold increase for Ohler7 on average). The other motifs did not affect or decreased the expression, maybe due to the disruption of sequences bound by unknown proteins. Overall, these results demonstrate positive effects on expression of most computationally-derived motif consensus sequences (except the repressive TTGTTrev).

**Figure 5.**
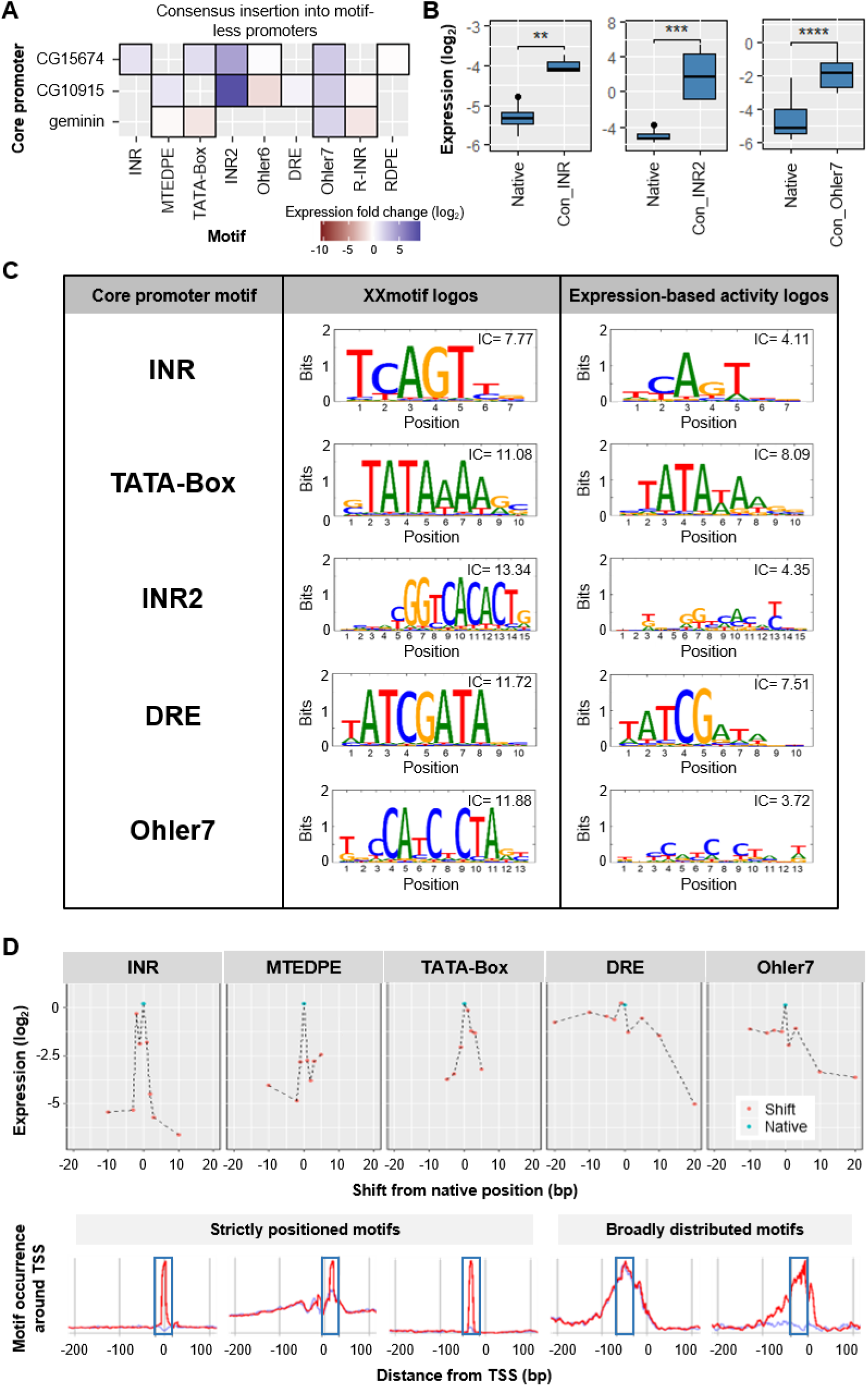
Consensus insertion, point mutation and positional shift. (A) Heatmap depicting the mean expression fold changes compared to wild-type expressions after replacing consensus insertion into motif-less core promoters. (B) Boxplots depicting log expression change and significance level upon inserting consensus motifs of INR, INR3 and Ohler7 motifs (columns in A) into the core promoters (rows in A). Left panel: INR into CG15674 (two-sample t-test *p =* 0.0033); middle panel: INR2 into CG10915 and CG15674 (Wilcoxon rank-sum test *p =* 0.00018); right panel: Ohler7 into Geminin, CG10915 and CG15674 (Wilcoxon rank-sum test *p =* 3.4*10^-5^). (C) Comparison of the *XXmotif* logos with the expression-based activity logos for INR, TATA-Box, INR2, DRE and Ohler7. Expression logos show an overall lower specificity. IC, information content. (D) Effect of motif positional shifts. Upper panel: log_2_ expression of native promoters (cyan dots) and promoters with motifs shifted relative to their original locations (red dots), for INR, MTEDPE, TATA-Box in cas, and DRE, Ohler7 in RpL36A. Lower panel: Motif occurrence around TSS (at position 0) discovered in the genome-wide analysis by *XXmotif.* The blue rectangular boxes indicate the −20 to 20 bp region surrounding the original positions of the motifs in the tested core promoters (strictly positioned INR, MTEDPE, TATA-Box in cas; broadly distributed DRE, Ohler7 in cas).

### Systematic point mutations enable the generation of expression-based PPMs and activity logos for core promoter motifs

We then systematically measured the influence on expression under all possible single base pair mutations of the motif consensus for various native promoters (details in **MATERIAL AND METHODS**). We recovered nearly all of the variants for the motifs INR, TATA-Box, INR2, DRE and Ohler7. In most cases the consensus sequences gave the highest expressions. Based on these expression measurements, we generated PPMs, and thereby activity logos for these motifs, which we compared with their *XXmotif* sequence-based logos (Figure 5C). For all motifs the expression consensus is identical to the computational one. All the expression-based activity logos are less specific, as indicated by their lower information content IC (Figure 5C, upper right corners) compared to those found in silico by *XXmotif.* An exception were the CG nucleotides in the DRE motif that have higher information content than the equivalent positions in the motif generated by *XXmotif,* suggesting their function as the primary recognition site for DREF binding.

In summary, although the computationally identified overrepresented sequence generally represents the best motif, the specificity of the individual nucleotides in the sequence tends to be overestimated. This is not surprising since computational motifs are derived based on over representation in promoter subgroups, which induce a bias towards higher specificity to distinguish them. In addition, their specificities is tuned by how strict the method is in accepting weak-strength motifs as true binding sites.

The positionally or functionally equivalent core promoter motifs from other architectures can hardly function as endogenous sequences While checking the features of core promoter motifs discovered by *XXmotif* (**Table S1**), we confirmed that certain motifs tend to locate in different core promoters within a similar region relative to TSS (like DRE and Ohler6 at around −100 to −7), or they share similar sequence features such as the “CA”s in INR, INR2 and Ohler7. We investigated whether positionally or functionally equivalent motifs (i.e. leading to similar decrease of expression after knockout) from other architectures could rescue the expression from knockouts. Three motif groups were tested: INR - INR2 - Ohler7 - R-INR; TATA-Box - Ohler6 - DRE; MTEDPE - RDPE.

For most of the motifs, we found that substitution could not recover the promoter activity, that is, substitution would yield the same or an only slightly higher expression than if the motif was knocked out (**Figure S5A**). An exception was the INR2, which could almost compensate for a INR knockout - showing a rescue effect (**Figure S5A**, upper left panel and **Figure S5B**; Wilcoxon rank-sum test *p =* 0.17 between the native expression and the INR2-substituted expression). Conversely, INR was not able to compensate for the loss of INR2 (**Figure S5A**, upper left panel). They both generally increased expression level compared to the native arrangement when substituting R-INR. This is likely due to the low intrinsic expression levels of R-INR-containing promoters, and matches the lack of R-INR knockout effect.

### Precise positioning of motifs is an essential feature of core promoter function

The *XXmotif* analysis showed the strong positional preferences of some motifs (**Table S1**). To test the functional relevance, we shifted the motifs around their native positions and checked the consequences on expressions.

Overall, varying motif positions from their position in the examined native promoters decreased the expression level, regardless of the shift direction (Figure 5D, upper panels). Additionally, the decrease in expression level correlated with the shift size. In the case of strongly positioned motifs (INR, MTEDPE and TATA-box), even small shifts (< 5 bp) led to a severe loss of expression, while less well-positioned motifs (DRE, Ohler7) showed milder effects when shifted (Figure 5D, upper panels). These position dependent expression patterns showed similar shapes as the genomic motif distribution within ±20 bp region of the most enriched motif locations (Figure 5D, lower panels).

In conclusion, the motif position is essential for core promoter function, because shifting affects the expression. Even single bp shifts can have strong effects. The genomic distributions of a motif reflect its measured expression pattern.

### A linear combination of individual motif features can largely explain the core promoter activity

Our results obtained from the pairwise knockout of motifs revealed the existence of superadditive or subadditive effects of individual motif features (Figure 4E - 4F and **S4B - S4G**). This prompted us to investigate how much of the expression level can be explained by the pure additive contributions of each motif feature. Therefore, we tested promoters combining all types of combinatorial mutations (varying motif strength, shift, and replacement) given a core promoter architecture (termed intra-architecture mutations; Figure 3B). We applied a linear regression analysis to predict log2 expression, assigning the covariate variables in the model as the qualitative indicators (0/1) of the individual mutation existence (**MATERIAL AND METHODS**). We obtained an average correlation of 88% (6 promoters tested) between predicted and experimentally measured log2 expression levels (Figure 6A and **S6A**). The coefficients learned by the models also correlate with expression levels of single mutation promoter (average correlation *PCC r =* 0.93; **Figure S6B**).

**Figure 6.**
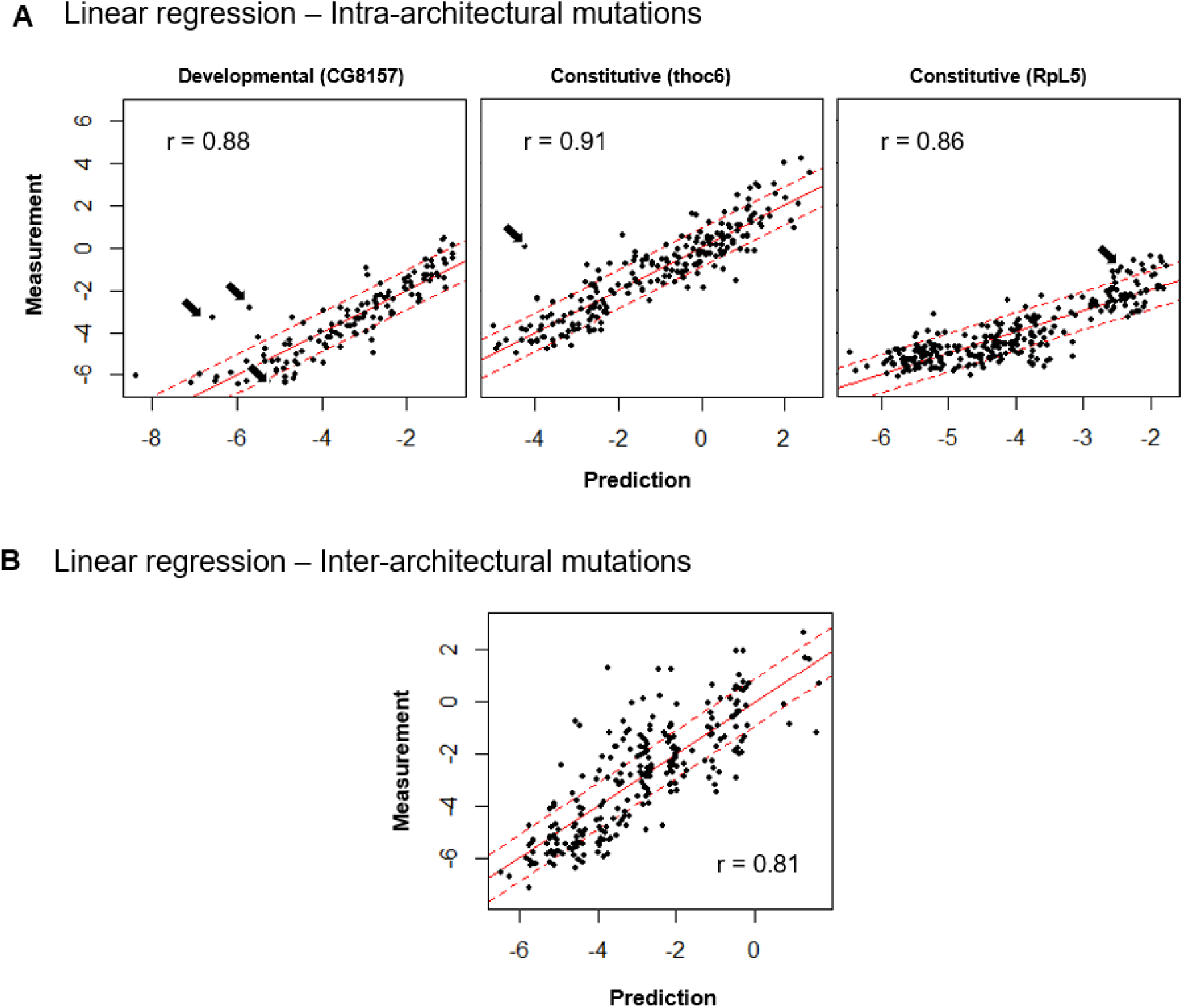
Linear regression modeling. (A) Linear regression applied to predict the synthetic promoter activity based on individual motif features (intra-architectural mutations). The measured expressions (on the y-axis) for 6 tested core promoter sequences with combinatorial motif mutations compared to the predicted expressions (on the x-axis) from the linear regression (log2 scale). Red solid line: y = x; red dashed lines: y = x ± 3×SD, where SD denotes the median of all standard deviations over all measured synthetic promoter constructs. It is an estimate for the noise in the expression measurements. The linear regression model can explain on average 88% of the variance in expression (average r = 0.88). (B) Linear regression analysis for inter-architectural block-wise combinatorial mutations. The measured expressions (on the y-axis) for inter-architectural block-wise combinatorial mutations compared to the predicted expressions (on the x-axis) from the linear regression fit.

As a more direct test without any fitting procedure, we also built an additive model to predict the activity of a given promoter with the intra-architectural combinatorial mutations based directly on the measurements of individual motif mutations (Figure S6C). The contribution of each feature (both motif strength and position) was assumed to be additive and was derived from the deviation between the corresponding motif-mutated sample compared to the native expression. Except for one promoter (cas, for which multiple single mutation constructs were not recovered during the cloning procedure; **MATERIAL AND METHODS**), we obtained a comparable mean correlation of 84%. To conclude, our results suggest the activity of a given synthetic core promoter is largely predicted from the linear combination of individual motif features. Both a linear regression model and a parameter-free additive model can explain most of the variance in expression. However, deviations are still observed, revealing the complex interplay between the factors involved. interplay between the factors involved.

### Motif context in core promoters influences expression

In addition to mutations applied to sequence motifs, we also tested the influence of the motif context on the expression level, that is, the sequence environment surrounding the motifs in the core promoter region (Figure 3C).

We first created promoter variants where either all motifs or motif contexts were shifted together, thus, maintaining the relative spacing of motifs while altering the sequence background in which they are located. In general, both cases led to the loss of expression; the effects were comparable or lower than those obtained from individual motif shifts (Figure S4I).

Besides the mutations applied within each native core promoter architecture, we also exchanged context sequences surrounding the motifs of a given promoter with foreign context sequences originating from other promoter architectures. The analysis revealed that overall the motifs preferred their native contexts (Figure S4J). For instance, the motifs from RpL5 resulted in an average more than 10-fold reduction of the expression levels when added into any of the other promoter contexts. When inserting motifs from any of the tested promoter architectures into motif-less core promoters (CG10915 and CG15674), they drastically improved the expression with a maximum increase of more than 55-folds (Figure S4J, blue squares). When comparing the obtained results with the wild-type expressions of the motif-origin promoters, the context from CG15674 could recover or even increase the expression of developmental promoters with their native motifs (∼ 25% expression increase for cas and > 2-fold increase for CG8157). Similarly, the context from the motif-less core promoter CG10915 could constitute a better promoter compared to the native Thoc6 (a constitutive promoter; with a ∼ 2.5-fold increase). Note that, although we checked if the various context effects could be explained by the classification as narrow peak (NP) or broad peak (BP) promoters, we did not see a clear relationship.

Given the effects observed for motif contexts and the strong predictability of core promoter activity based on individual motifs, we wondered which role play the context sequences surrounding the motifs in defining core promoter function (inter-architectural mutations, Figure 3D). Similarly to the previous section, we applied a linear regression model on the results obtained with the inter-architectural block-wise combinatorial mutations (Figure 6B; details in Supplementary Discussion 2). Remarkably, the predicted values showed here too a good correlation with the measured expressions (PCC r = 0.81, p < 2.2×10-16), recapitulating the possible additivity for sequence blocks even among various promoter architectures.

To summarize, our results show that the motifs do not contain all the information. The context sequences surrounding the motifs in core promoters also play an important role in defining the activity. These effects are however generally less prominent. The block sections which contain motifs together with their surrounding context sequences largely function in a linear way for setting expression levels.

### Ecdysone responsiveness correlates with the core promoter architecture

Finally, we checked the global ecdysone responsiveness for our entire synthetic promoter library. An *a priori* scenario was the possible repressor role of unliganded EcR known before (Cherbas et al., 1991; Dobens et al., 1991). We first performed control experiments which confirmed that the activity of a synthetic promoter containing the *block 2* sequence without induction was similar to the activity of the same core promoter sequence but without EcR/USP binding sites in *block 2* (data not shown). This suggested that the measurements without ecdysone induction in our experiments represent the basal activity of the tested synthetic promoters.

Ecdysone activation increased the expression of almost all promoter candidates (both native and mutated) tested in our experiments. The ecdysone responsiveness spanned a range of 1000-fold difference between the highest and lowest effect. We found (Figure 7A - 7B and S7A - S7C) that developmental core promoters (green dots in Figure S7A) were highly induced with an average > 20-fold activity increase, while constitutive core promoters (red dots in Figure S7A) showed much weaker responses (around a 4-fold increase on average; Figure S7B). Given ecdysone is a developmental stimulus, it should be expected to preferably activate developmental core promoters. Some housekeeping core promoters with already high basal expression levels without ecdysone stimulation (log_2_ expressions > 2; on the right of the red dotted line in Figure 7A) exhibited much smaller activations, suggesting saturation of promoter expression level that cannot be further enhanced.

**Figure 7.**
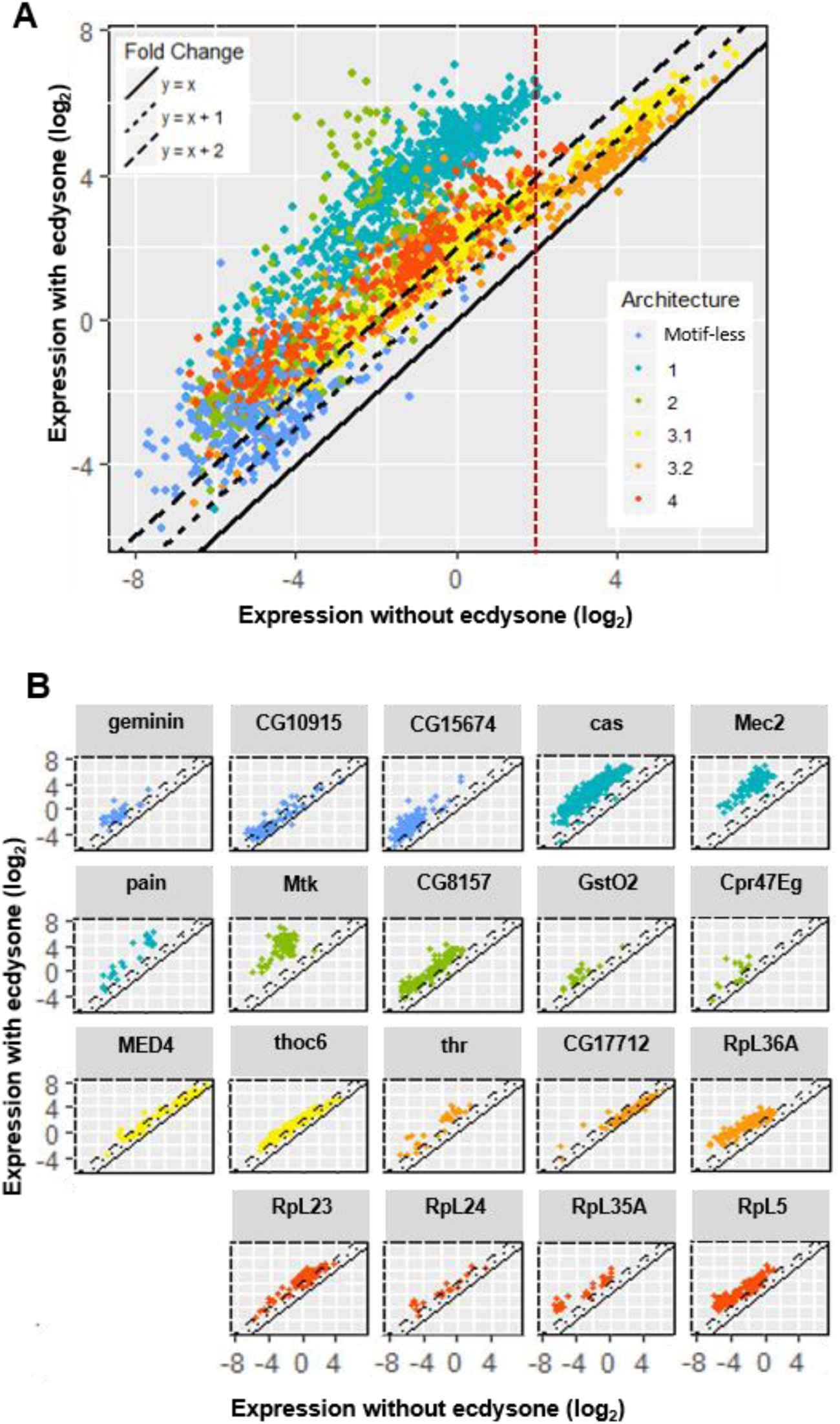
Ecdysone inducibility. (A) Scatterplot depicting the expression measurements with ecdysone induction versus measurements without ecdysone for all tested promoters separated by promoter architecture. Each color represents one architecture (color-code indicated in the insert). Three types of line are used to indicate the expression fold change with no increase (y = x; solid line), 2-fold increase (y = x + 1; dotted line) and 4-fold increase (y = x + 2; dashed line). Red vertical dashed line: log_2_ basal expressions = 2. (B) Comparison of the expression fold changes versus measurement values without ecdysone for all native promoters (grouped by native core promoter sequences). The colors refer to different core promoter architectures. Black line: linear regression (with the 95% confidence interval shown in gray, *PCC r* and *p* are shown for each group).

To gain deeper insight, we checked the ecdyson responsiveness of each promoter individually (Figure S7C). Here, the ecdysone responsiveness is defined as the ratio between the induced and uninduced expression level; also referred to as the ecdysone inducibility or the expression fold change caused by the ecdysone induction. We found a generally negative correlation between inducibility and expression level without ecdysone stimulation (Figure S7C; the only exception was a group of sequences derived from pain core promoter, which had increased inducibility with higher expression; *r =* 0.51; *p =* 0.012 s.) The higher the expression level, the lower the inducibility, which is consistent with the low activation measured for promoters with high basal expression level. The negative correlation was more significant for constitutive core promoters than developmental ones (Wilcoxon rank-sum test *p =* 0.0054; Figure S7B).

The ecdysone inducibility was generally independent of nearly all single motif knockout mutations (Figure S7D) with an exception of INR (a slightly negative effect of ∼ 2.3-fold reduction on average, Wilcoxon rank-sum test *p =* 2.1×10-5). Similarly, the motif consensus sequences didn’t dramatically affect the ecdysone responsiveness (< 20% reduction on average; Figure S7E).

Together, our results demonstrate a correlation between the ecdysone responsiveness and the core promoter architecture. Ecdysone can induce both developmental and constitutive core promoters but drives higher stimulations on developmental ones. The ecdysone inducibility generally decreases with the expression level for a given promoter: the higher the activity, the more difficult it seems to be to boost further expression level. Very strong promoters are barely inducible, probably due to promoter activity saturation. Finally, motif disruption has only minor influence on the ecdysone responsiveness of the core promoter.

## DISCUSSION

Our results reinforce the conclusions drawn from other smaller-scale studies for the roles of core promoter motifs in determining transcriptional output, also generalizing their effects to more promoter architectures. However, the major contribution of this work is to bring new insights into *D. mel.* core promoter function.

We demonstrate that the well-known functional motifs like INR, TATA-Box, MTEDPE, INR2 (more widely known as Ohler1 or motif 1), DRE and Ohler7 are crucial for gene expression. Their roles are unique and they cannot be replaced by positionally or functionally similar motifs from other architectures. Pairwise knockouts mostly elicit more significantly negative effects on transcription, and these effects show in some cases superadditivity. Conversely, most of the motif consensus sequences tend to increase core promoter activity. All these findings are consistent between different core promoters, and emphasizes again the importance of the sequence motifs for core promoter function.

However, not all well-characterized motifs have a significant effect on expression. This is especially the case with R-INR (more widely known as the TCT motif), which is also the only CA less TSS-motif and is part of a specialized TCT-based Pol II transcription system, distinct from the INR-based system (Parry et al., 2010a). This particularity might explain why this motif is special in the sense that it surprisingly makes almost no contribution to the expression although it exists in nearly all ribosomal protein gene promoters in *D. mel.* In contrast, housekeeping core promoter motifs like INR2 and Ohler6 that co-occur in multiple promoters show stronger influence in our data. It is known that more than half of the ribosomal core promoters contain this INR2 motif (Ma et al., 2009). A recent study proposed that the INR2 binding protein M1BP can act as an intermediary factor to recruit TRF2 for proper transcription of ribosomal protein genes (Baumann and Gilmour, 2017). Our perturbation analysis of INR2 in various ribosomal promoter backgrounds supports their finding. The results we obtained with Ohler6 also suggest that the unknown TF(s) that bind to it may function similarly as M1BP.

Among the four tested novel motif candidates discovered by *XXmotif,* we identified TTGTTrev and RDPE as having measurable effects on expression after mutation, hereby confirming their biological relevance. TTGTTrev shares a similar function with a negative regulatory element for binding of a transcriptional repressor AEF-1. The occurrence of RDPE is highly correlated with R-INR and can partially replace the function of MTEDPE in developmental architectures. However, we note that the mutations in the two newly discovered motifs like TTGTT and CGpal show little effect on expression, suggesting these two computationally derived over-represented sequences lack functional importance as core promoter elements. They are therefore likely to represent binding sites of transcription factors that are not expressed in our experiments. Due to the similarity of TTGTT with R-INR, this motif may act as a redundant version of the R-INR motif.

Our highly sensitive assay can also accurately capture the partially subtle expression changes caused by single base pair variations of motifs. We confirm that the most-overrepresented sequence of a given motif in the genome mainly stands for its best functional form, but we also saw differences with the computationally derived matrices: our expression-based activity logos are generally less specific. The two kinds of motifs are complementary since they reflect different phenomena: *in silico* discovered motifs are expected to reflect binding affinities, whereas the expression measurements measure effect on transcription initiation, which could be buffered, for example, by alternative pathways / coactivator complexes.

Altering motif positions overall decreases expression. Several studies have suggested the exact spacing is essential for synergism between the core promoter motifs to function as active pairs to recruit GTFs along with Pol II for accurate transcription initiation (Burke and Kadonaga, 1997; Emami et al., 1997; Gershenzon and Ioshikhes, 2005; Gershenzon et al., 2006; O’Shea-Greenfield and Smale, 1992). Our results are in line with these previous findings for strictly positioned motifs such as INR, MTEDPE and TATA-Box. Their locations and spacings are highly restricted for the effective binding of the TFIID to nucleate the PIC. Other motifs that can function over wide ranges and are not necessary for constituting the major machinery, e.g., DRE, Ohler6 and Ohler7, show less stringent location requirement and smaller effects on expression, as long as they do not disrupt other sequence features.

Importantly, we also demonstrate that not only the core promoter motifs but also their context sequences determine expression output, giving insights into the debated role of motif flanking and context sequences of core promoters. Our results uncover that sequence motifs mostly prefer their native context. Remarkably, although only INR and INR-like motifs including INR2 and Ohler7 can drive higher expression when their consensus sequences are inserted into motif-less core promoters, the motif combinations from almost all the other defined architectures can result in a substantial increase of expression level, revealing the importance of motif synergism. We also see an influence of the sequence context independent of the motifs, which may obey complicated rules. It is however beyond the scope of this study.

Considering that pairwise motif disruption already suggests certain levels of synergistic effects, the higher-order combinatorial effect of mutant motifs and their context on expression may be more difficult to understand. To dissect this complexity of the mutant combinations, we used a linear regression model to check how much of the core promoter activity can be correlated with individual effects. To our surprise, we found that the expression changes caused by single mutations of sequence motifs joined in a linear fashion can largely predict the output of the free mutant combinations. Hence, promoter expression levels of mixed and combined motifs can largely be explained by simple linear addition of their individual contributions. We also extended the sequence features from simply the motifs alone to larger sequence blocks that contain motifs together with their context surrounding sequences. We also found that a linear model describes the expression of these inter-architectural block combinations well. A linear combination of individual sequence features like the motifs or wider sequence blocks including their context sequences can account for two-thirds of the variance in expression levels as regulated by the core promoter. To unravel the nonlinear interactions more mutational data and more detailed models will be necessary.

The ecdysone responsiveness highly correlates with core promoter architecture. This developmental stimulus functions more strongly on developmental core promoters. There is a generally negative correlation between the ecdysone responsiveness and the basal expression level. Our strongest promoters can barely be induced by ecdysone. The higher the expression level, the more difficult it is to further boost the signal, hinting at the saturation of the promoter expression. This effect is stronger for constitutive core promoters, showing their less efficient activation. The disruption of INR in developmental core promoters can lead to a reduction in the ecdysone responsiveness, which is consistent with what was reported in a previous study in *Spodoptera frugiperda* (Jones et al., 2012). Taken together, the different sequence motifs composing distinct core promoter architectures can predict their ecdysone responsiveness: developmental core promoters can get much higher induction.

Finally, by investigating the effect of potential nucleosome binding, we observe moderate effects on expression (compared to motif knockouts) driven by these different potential nucleosomal backgrounds. Note that although we checked nucleosomal presence on plasmid for one construct, it is not known if our promoters have native nucleosome occupancy. We however find greater expression variation for housekeeping and ribosomal core promoters than developmental core promoters when changing the TSS nucleosomal sequence downstream the TSS (block 7); this suggests the significance of the genomic +1 nucleosomal sequences for the function of constitutive core promoters.

## ACKNOWLEDGMENTS

We thank Prof. Dr. Achim Tresch for fruitful discussion in the initiation phase of the project and Prof. Patrick Cramer and Prof. Roland Beckmann for their support.

## AUTHOR CONTRIBUTIONS

Z.Q. carried out most of the experimental work and analyzed the data with the help of P.B., C.L., M.M., A.R., J.P.M. and M.S.; M.H. and A.S.K carried out the computational analysis and designed the promoter sequences; C.J. designed the experiments and analyzed the data; U.G., U.U. and J.S. designed the study; U.G., C.G., U.U., and J.S. supervised the work; Z.Q. and C.J. wrote the paper with contributions from the other authors.

## MATERIAL AND METHODS

### De novo motif search using the *XXmotif* algorithm

In a previous work, we devised the *XXmotif* (exhaustive evaluation of matrix motifs), a P-value-based regulatory motif discovery tool using position weight matrices (PWMs) (Hartmann et al., 2013; Luehr et al., 2012). In brief, we firstly defined 19 gene sets based on experimentally derived genome-wide features, including expression strengths and variations throughout developmental stages (Graveley et al., 2011), Pol II stalling (Hendrix et al., 2008; Zeitlinger et al., 2007) and TSSs mapping from CAGE data (Hoskins et al., 2011; Ni et al., 2010). We then applied *XXmotif* for the de novo motif search in the core promoter regions of these genes and were able to identify widely known motifs as well as some novel motif candidates with optimized PWMs based on enrichment, localization and conservation (Hartmann, 2012).

### Synthetic promoters design

#### Building blocks

We designed synthetic promoter constructs by dividing the promoter region into 7 building blocks (Figure 1A-B): *block 3-6* (131 bp in length) was the motif-rich core promoter region (−80 to +50 bp around the TSS) with native and mutated sequences from different core promoter architectures to investigate the effects of sequence motifs; *block 2* (73 bp) represented the EcREs, which contained the binding sites for the ecdysone receptors to recruit the steroid hormone ecdysone for transcriptional activation; *block 1* (239 bp) and *block 7* (240 bp) were used for testing the influence of nucleosomal sequence context. The entire lengths for the designed synthetic promoters inserted into the vector backbones were 703 bp with *block 7* and 459 bp without *block 7*.

#### Nucleosomal context (*block 1* and *block 7*)

After MNase digestion of chromatin, genome-wide nucleosome maps were generated (data not schown). 12 gene promoters were selected according to their pattern of nucleosome positioning and occupancy relative to their TSS (especially ±1 nucleosomes) and pairs of *block 1* and *block 7* sequences representing different potential ±1 nucleosome patterns were selected (sequences in **Table S5 and S6**). The block 1 and 7 sequences were synthesized either by PCR amplification from the genomic DNA (isolated from sequenced fly strain, stock number 2057 in Bloomington *Drosophila* Stock Center) or by oligo synthesis from Life Technologies (for HindIII recognition sites mutated and ATGs mutated sequences). All synthesized sequences of *block 1s* and *block 7s* contained BsaI sites and assembly overhangs, and they were stored in TOPO vectors (Zero Blunt TOPO PCR Cloning Kit, Invitrogen). In the experiments, we tested *block 1* and *block 7* in pairs with all 19 native core promoter *blocks 3-6,* five out of which were then selected to combine with all free combinations of *block 1* and *block 7* (one from each architecture with activities covering the entire dynamic range: CG15674 (motif-less), Mec2 (Ar.1), Mtk (Ar.2), CG17712 (Ar.3), RpL23 (Ar.4)). We also constructed synthetic promoters containing only *block 1* (without *block 7*) for these five wild-type *blocks 3-6.* One pair of *block* 1.11 and *block* 7.11 was selected based on its high expression level and used as the fixed nucleosomal sequence context for highly mutated blocks 3-6.

#### Ecdysone receptor binding site (*block 2*)

The *block 2,* which contained three EcR/USP heterodimer binding sites with 17 bp spacers in between, was synthesized by oligo annealing (5’-gc<u>GGTCTC</u>AATGA<u>agttcattgacct</u>agtgag aattcaca<u>gcgagttcattgacc</u>tactcaaggcatacatga<u>agttcattgacct</u>GGATTGAGACCgc-3’; lowercase with underline: EcR/USP binding sites from JASPAR database (Khan et al., 2018); italic: assembly overhangs; uppercase with underline: BsaI restriction sites). Selection of the native core promoter set (*blocks 3-6*) From the four core promoter architectures (including two subclasses Ar.3.1 and Ar.3.2 of the housekeeping Ar.3) and one additional architecture without having any known motif termed motif-less promoters, we chose 2-4 native core promoters each with high or intermediate to low expressions according to their maximum expression levels in S2 cells (previous RNA-seq data generated by our group; position −80 to +50 relative to TSS which was set to be position 0; block 3: −80 to −35, block 4: −34 to −10, block 5: −9 to +8, block 6: +9 to +50). In total, we thus selected 19 wild-type core promoters, some of which have mixed architectures due to different motifs co-occurrence (Figure 2; their 131 nt sequences listed in **Table S3**). The annotation of core promoter motifs in these sequences was carried out by motif search using *XXmotif* according to previously defined motif features (summarized in **Table S1**). In addition, we mutated TSS downstream ATGs in the original sequences to TAGs to remove unwanted translation start sites.

### Mutation with different strengths of motifs

Various kinds of mutations were designed for these native core promoters, including mutations for motifs within each core promoter (main mutations shown in Figure 3) and block-wise mutations between different core promoters. We scanned every designed sequence with our PWMs to check if the mutants we created would lead to undesirable side mutational effects, e.g., the creation of new motifs/TF binding sites or disruption of other motifs (as those unintended mutations would cause expression changes).

#### Knockout of motifs

For knocking out individual motifs in 16 native core promoters (excluding three motiless promoter sequences), two versions of sequences were used as substitutions: random sequences and background sequences. Random sequences were generated by sampling sequences having the same length with the target motifs and checking with the *XXmotif* derived motif list to make sure no known core promoter motif inside (whose PWM scores lower than the threshold, threshold score of each motif listed in **Table S2**). These random sequences were not fixed for the same motif in different promoters (every random sequence was different). Background sequence was a fixed sequence from the identical position of the target motif in the motif-less core promoter CG15674 (due to the various positions of a certain motif in different promoters, the background sequence might vary). Knockout of all motifs in a given promoter was designed in the same way, using both random and background sequences. Pairwise knockout of motifs only used random sequences for replacing two original motifs at the same time.

#### Consensus replacement of motifs

For the nine main motifs INR, MTEDPE, TATA-Box, INR2, Ohler6, DRE, Ohler7, R-INR and RDPE, we replaced them in native core promoters with the consensus sequences derived from *XXmotif.* Additionally, these consensus sequences were also inserted into the three motif-less core promoters with their start positions at the peaks of the native motif distribution (motif distribution shown in the column “Distribution” of **Table S1**).

#### Replacing native motifs with their alternatives of various strengths

Alternatives with different PWM scores for the nine main motifs mentioned above were randomly generated, making sure that their scores either evenly covered several score bins below the threshold and the maximum.

#### Point mutation of motifs

For the 12 motifs INR, MTEDPE, CGpal, TATA-Box, INR2, Ohler6, DRE, Ohler7, R-INR, RDPE, TTGTT and TTGTTrev, we designed all possible single base pair mutations around the motif’s consensus sequence. This was done for each motif within a selected native core promoter configuration: INR in Mec2; MTEDPE and CGpal in Cas; TATA-Box in CG8157; INR2, Ohler6 and TTGTTrev in Thoc6; DRE, Ohler7 and TTGTT in RpL36A; R-INR and RDPE in RpL5. Additionally, INR, DRE, Ohler7 and R-INR were also checked in an motif-less context sequence obtained from CG10915, with the insertion of each consensus sequence.

#### Substitution of motifs

The target motif was firstly knocked out with a random sequence. The motif sequence for substitution was also randomly sampled with a PWM score above the threshold and was always the same for each motif. Three combinations were tested here: INR (7 nt) - INR2 (15 nt) - Ohler7 (13 nt) - R-INR (11 nt); TATA-Box (10 nt) - Ohler6 (10 nt) - DRE (10 nt); MTEDPE (17 nt) - RDPE (17 nt). For INR-like motifs with various lengths, the supposed position for TSS (3rd position in INR, 10th in INR2, 5th in Ohler7 and 6th in R-INR; based on the motif start positions listed in **Table S1**) was aligned when replacing the sequence.

#### Positional shift of motifs

Positional shifts were designed for individual motifs and all motifs together in a given core promoter, as well as for sequence context surrounding motifs (motifs kept at the original positions). For strictly positioned motifs like INR, MTEDPE and TATA-Box, shifts of 1, 2, 3, 5, 10 bp either downstream or upstream were applied; for less well-positioned housekeeping core promoter motifs like DRE and Ohler7, larger distances were chosen (±1, ±3, ±5, ±10, ±20 bp).

#### Other combinatorial mutations

Further combinatorial mutations were designed to the motif-rich core region, including free combinations of mutations both within defined core promoter architectures and between them (termed as intra-architectural motif-wise and inter-architectural block-wise combinatorial mutations). In addition, context sequences surrounding the motifs were also tested by exchanging them between different core promoters.

For testing these combinatorial mutations, one representative core promoter sequence from each architecture with motifs located within distinct block regions was selected: Cas (Ar.1), CG8157 (Ar.2), Thoc6 (Ar.3.1), RpL36AN (Ar.3.2) and RpL5 (Ar.4). The synthetic promoter RpL36AN was derived from the native RpL36A (Ar.3.2) shifting the TSS position 16 nt upstream in order to shift all motifs into the blocks where they occur most frequently, based on the distributions generated by XXmotif. In addition to the five core promoter sequences tested systematically in all three types of combinatorial mutations, several other native sequences were also included (MED4 for intra-architectural mutations; Mtk and Cpr47Eg for inter-architectural mutations; CG10915 and CG15674 for context exchange).

#### Intra-architectural motif-wise combinatorial mutations

Multiple motif-wise mutations for altering both motif strength and motif position within a core promoter sequence were performed here. The MED4 (Ar.3.1) was selected because of its strong native activity level, which ensures a relatively strong luminescence signal even after severe combinatorial mutations. Single mutations (knockouts, replacing by the consensus or alternatives with different PWM scores and positional shifts) for individual motifs in each core promoter were re-designed in the same way as described before but kept the same in all intra-architectural combinatorial mutations. Shifts of motifs were made within shorter ranges (±1 bp or ±5 bp).

#### Inter-architectural block-wise combinatorial mutations

We applied block-wise swaps between different core promoter sequences here. Two additional sequences Mtk and Cpr47Eg were included to provide extra block patterns. In detail, block pieces from 7 native core promoters were selected and freely combined to construct the synthetic *block 3-6* regions: four block 3s from CG8157 (background sequence of Ar.2), RpL36AN (background sequence of Ar.3.2, BP), RpL5 (Ohler6 existed), Cpr47Eg (CGpal existed); five block 4s from Cas, CG8157, Thoc6, RpL36AN, RpL5; four block 5s from CG8157, Thoc6, RpL36AN, RpL5; six block 6s from Cas, CG8157, Thoc6, RpL36AN, RpL5, Mtk (background sequences of Ar.2).

#### Context exchange

All motifs in a given core promoter were knocked out using the same sequences designed for single knockouts in intra-architectural combinatorial mutations. All motifs from other core promoter sequences were inserted into this context at their native positions (Figure 3C). Two motif-less core promoter contexts were also included: CG10915 (BP) and CG15674 (Adams et al.).

### Experimental setup and procedures

#### Reporter and control plasmids for dual luciferase assay

A two-vector system was used in the experiments. Firefly reporter vector backbone was derived from a commercial vector pGL4.13 with luc2 firefly gene (Promega). HindIII and BglII restriction enzymes (Khan et al.) were used to cut out the SV40 early enhancer/promoter region in the original plasmid. To insert BsaI sites and 4 bp overhangs, two dsDNAs with HindIII and BglII sites were generated by oligo annealing: for the constructs containing a *block 7* (sequences listed in **Table S6**), the following sequence was used: gcagatctgcGAACTGAGACCgtcgacgcaaggcctgcaattaatgcagcggccgatcggcatatgGGTC TCA CCACcaaagcttcg (only forward sequence; BglII or HindIII restriction sites: lowercase with underline; overhangs: italic; BsaI restriction sites: uppercase with underline); the sequence used for the constructs without *block 7* was: gcagatctgcGAACTGAGACCgtcgacgcaaggcctgca attaatgcagcggccgatcggcatatgGGTCTCATCTGcaaagcttcg. After enzymes digestion and gel purification (QIAquick Gel Extraction Kit, Qiagen) of both vector and inserted DNAs, ligation (Rapid DNA Ligation Kit, Roche) was performed to obtain the two final vector backbones (4299 bp), named as BB0 for the constructs without *block 7* and BB1 for the constructs containing a *block 7.*

Renilla control plasmid (3630 bp) was derived from another commercial vector pGL4.70 with the hRluc renilla gene (Promega) by insertion of a moderate-strength P transposase (pTran) promoter between NheI and XhoI sites. The pTran promoter was cloned from a vector created in the lab pKF1 (derived from a P-element sequence, position 34-141 according to (O’Hare & Rubin, 1983)) using primers: 5’-GC<u>GCTAGC</u>AGCCGAAGCTTACCGAAGTATAC-3’, 5’-GC<u>CTCGAG</u>CCACGTAAGGGTTAATGTTTTC-3’ (underlines: NheI and XhoI restriction sites).

Several inter-plate controls were used in the experiments. The negative control was one commercial vector pUC19 (Khan et al.). There were two positive controls: one was pGL4.10 vector (Promega, with luc2 firefly gene) with pTran promoter inserted between NheI and XhoI sites, termed as pUG9, whose signal was used in data normalization procedure (4350 bp); the other one was a synthetic test plasmid pZQ3 (4691 bp) with moderate promoter activity which contains our firefly reporter backbone BB0 and blocks 1-6 for ecdysone inducibility check: *Block 1.3* (all *block 1* sequences listed in **Table S5**) + *Block 2* (sequence in Section 4.2) + *Block 3-6* with INR and DPE motifs (sequence: GGCTCCGAATTCGCCCTTTTCCCAGGGCGGCAGAGGCAAAAATTTGCCGA TCCCAGAGCCAGCCGACTCATTCAAAGCTCCGACTTCGTTGCGTGCACACAGA GTCTCAAGGGCGACCCAGCTTT).

#### Cloning

For carrying out our large-scale systematic analysis, we developed a high-throughput experimental pipeline using automated robotic systems (Figure S1). After preparation of each construct block (*block 1* and *block 7*: PCR amplification from the fly genome or oligo synthesis; *block 2*: oligo annealing; *block 3-6*: PCR amplification from the synthetic library (Agilent Technologies) according to mutation families), Golden Gate cloning (BsaI cloning) was applied to join them with the vector backbones sequentially. Then, the newly synthesized reporter plasmids were transformed into electrocompetent *E. coli*, followed by plating bacteria on one-well plates, this way facilitating automated colonies picking using the robotic workstation. After bacterial growth in 48-well plates, we rearranged them into 96-well LB plates and prepared the library for next-generation sequencing with two-step PCR using nested barcode primers. Based on the sequencing results, replicates and bad clones were screened out and DNAs from confirmed positive clones were isolated. These firefly reporter plasmids containing all the distinct promoters were then used for transient co-transfection into *D. mel.* S2 cells together with the renilla control plasmid in 96-well plates. After overnight incubation, cells were treated with ecdysone for another 2 hours. Four cell culture 96-wellplates were pulled into 384-well plates for the final dual luciferase assay readout in order to use less substrate for the luciferase assays.

#### Automation

We used two independent robot platforms with a similar basic configuration of pipettor systems (Biomek NXP automated workstations with Multichannel-96 and Span-8 pipetting model, Beckman Coulter). Additional instruments were integrated with the original workstations including incubators (Incubator Shaker DWP, Inheco), thermocyclers (Biometra TRobot, Analytik Jena), barcode printer (Microplate Print & Apply, Beckman Coulter), barcode reader (Compact Laser Barcode Scanner, Omron Microscan), plate reader (SpectraMax Paradigm Multi-Mode Microplate Reader, Molecular Devices), plate sealer (Wasp, Kbiosystems). They were designed for maximum flexibility to perform many different experiments. Specifically, one system is dedicated to bacterial experiments, mainly the cloning-related work: colony picking, colony PCR, hitpicking for positive clones, DNA isolation and concentration measurement. The colony picking is a customized feature of this robotic configuration. The other system is dedicated to *Drosophila* cell assays: transient co-transfection, ecdysone treatment and luciferase assay readout. In addition, an electronic multichannel pipette on an assistant robot (VIAFLO Electronic Multichannel Pipette + ASSIST Pipetting Robot, INTEGRA) was used for automated cell plating into 96-well plates.

#### Synthetic library amplification

*Block 3-6*s for the motif-rich core regions of our synthetic promoter constructs were amplified from a library synthesized by Agilent Technologies (LeProust et al., 2010) together with BsaI sites, relevant overhangs and unique primer sequences referred to distinct mutation families, in total 3826 fully designed oligonucleotides (in total ∼ 200 nt long for each sequence). The entire oligo pool (lyophilized, 10 pmol) was dissolved in 100 μ! Elution buffer (Qiagen) and shaken at room temperature (RT) for 30 min at 450 rpm and 10 min at 950 rpm. 0.5 μl of library DNA was used to amplify the specific sequence family (native sequences or one of distinct mutation families) in a 20 μl PCR reaction, which also included 1.25 μl of both forward and reverse 10 μΜ customized primers, 4 μl 5* Herculase II reaction buffer, 0.5 μl 10 mM dNTP mix and 0.5 μl Herculase II fusion DNA polymerase (Agilent Technologies). PCR parameters were as follows: 98 °C for 3 min; followed by 15 cycles of 98 °C for 80 s, 54 °C for 30 s, 72 °C for 40 s; 72 °C for 10 min. Each PCR reaction was purified with the QIAquick PCR purification kit (Qiagen) according to the manufacturer’s instructions and eluted in 30 μl of nuclease-free water (Qiagen).

#### Golden Gate cloning and transformation

BsaI restriction enzyme (10,000 U/ml, NEB) and T4 DNA ligase (3 U/μί, Promega) were applied to assemble all of the synthetic promoter blocks sequentially and simultaneously into the firefly reporter vector backbone in a one-pot reaction. For each 20 μl reaction, DNA master mix contained equimolar amount (80 fmol) of each part: *block 1* in TOPO vector (3784 bp), *block 2* (99 bp), *block 3-6* (200 bp), *block 7* in TOPO vector (3785 bp, if needed) and backbone (4299 bp) together with 2 μl BsaI, 2 μl T4 DNA ligase and 2 μl 10* ligase buffer. The cloning protocol included 3 steps: (1) 20 cycles of 37 °C for 2 min, 16 °C for 3 min; followed by 50 °C for 5 min and 80 °C for 5 min; (2) After adding 1 μl BsaI, 1 μl T4 DNA ligase, 1 μl 10 mM ATP: 16 °C for 20 min; 15 cycles of 37 °C for 2 min, 16 °C for 3 min; followed by 50 °C for 5 min and 80 °C for 5 min; (3) After adding again 1 μΙ Bsal: 37 °C for 10 min, 50 °C for 20 min, 80 °C for 10 min and ramp down to 25 °C by 0.1 °C/s. After BsaI cloning, 2 μl of the reaction mix was transformed into 40 μl of electrocompetent TOP10 E. coli cells (homemade). After electroporation (1.8 kV for 0.1 cm cuvettes, Gene Pulser, Bio-Rad) and 1 ml SOC medium (homemade) addition, cells were incubated for 1 h at 37 °C (shaking at 450 rpm) and plated 100 μl onto prewarmed 1-well LB-agar plates supplemented with 100 μg/ml Ampicillin.

#### Colony picking

After overnight incubation at 37 °C, the 1-well plates were ready for colony picking. Span-8 pipetting system on the robot was used to automatically pick individual colonies (customized protocol) into two 48-well plates (Riplate SW 48, 5 ml, Riplate) with 2.4 ml LB-Ampicillin medium (Ampicillin concentration: 120 μg/ml). The plates were incubated for 16 h at 37 °C (horizontally shaking at 180 rpm) and rearranged into one 96-well plate (MegaBlock 96 Well, 2.2 ml, Sarstedt). 110 μl/well of bacteria was used to create glycerol stock plate (Round 96 Well Storage Plates, U-bottom, 330 μl, 4titude) and 30 μl/well for PCR plate (FrameStar 96 Well Skirted PCR Plate, 4titude) ready for sequencing library preparation. Since in the previous cloning step, the sequences from the same mutation family were all mixed together, it is technically impossible to recover all of them during the colony picking step. Therefore, we did over-picking of the colonies and were able to recover in total more than 3000 of the designed sequences.

#### Next-generation sequencing of the picked clones

Two-step PCR with nested barcode primers was implemented for library preparation. The forward and reverse primers for 1st PCR targeted the sequences in *block 2* and vector backbone respectively with specific barcodes (*block 1* was always known in the BsaI cloning procedure). 2 μl/well of bacteria were used to set up a 25 μl PCR reaction containing 1 μl homemade Taq/Pfu polymerase mix, 2.5 μl primer mix (forward and reverse each 500 nM), 1 μl 25 mM MgCl2, 2.5 Ml 10* buffer, 1 μl 2.5 mM dNTP. 96-well plate PCRs were performed in the thermocyclers integrated on the robot (96 °C for 7 min; 3 cycles of 94 °C for 30 s, 68 °C for 30 s, 72 °C for 2 min; followed by 3 cycles of 94 °C for 30 s, 64 °C for 30 s, 72 °C for 2 min; 17 cycles of 94 °C for 30 s, 56 °C for 30 s, 72 °C for 2 min). 5 μl/well of the product from each 1st PCR plate was pooled into one specific well of the collection plate (Deepwell plate 96/500 μ!, Eppendorf; each well containing all 96 samples from one 1st PCR plate). 3.5 μl/well was then used as template for 2nd PCR in a 50 μl reaction together with 0.5 μl Herculase II fusion DNA polymerase (Agilent Technologies), 10 μl 5* Herculase II reaction buffer, 1.25 μΙ 10 mM dNTP mix and 5 μΙ each of Illumina index primers (Nextera XT Index Kit v2, Index 1 (i7) Adapters and Index 2 (i5) Adapters, Illumina). So each well of 2nd PCR plate (each 1 st PCR plate samples) got a unique pair of index adapters. PCR was performed as the same protocol for 1st PCR. The final products were pooled and purified using Agencourt AMPure XP magnetic beads (Beckman Coulter) according to the manufacturer’s instructions. Next-generation sequencing (Illumina HiSeq1500) was performed by the LAFUGA sequencing facility at the Gene Center LMU Munich.

#### Hitpicking and DNA isolation

Automated hitpicking of positive clones from glycerol stock plates was carried out using our robotic system. 75 μΙ of the samples in the original plates were reformatted into the final 96-well glycerol stock plates (Round 96 Well Storage Plates, U-bottom, 330 μ× 4titude) and 20 μ! were used for reinoculation in 48-well plates (Riplate SW 48, 5ml, Riplate) with 2.4 ml LB-Ampicillin medium (Ampicillin concentration: 120 μg/ml). The plates were incubated for 17 h at 37 °C (horizontally shaking at 180 rpm) and rearranged into one 96-well plate (1.2 ml/well; MegaBlock 96 Well, 2.2ml, Sarstedt). After centrifugation at 5000 g for 15 min, the supernatant was discarded and cell pellets were stored at −20 °C ready for DNA isolation. Minipreps in 96-well plate format was performed with Wizard MagneSil TfxTM System (Promega) on the robotic workstation according to the manufacturer’s instructions. DNA concentrations were measured using the SpectraMax Microplate Reader integrated on the robot (5 μl DNA samples on the SpectraDrop Micro-Volume Microplates, Molecular Devices).

#### Cell culture

*Drosophila Megaloster* S2 cells were firstly thawed at passage 12 with Schneider’s Drosophila Medium (Bio&Sell, supplemented with 10% FBS (Fetal Bovine Serum, Biochrom)) and later cultivated in Express Five SFM medium (protein-free and serum-free, Invitrogen). One bottle of the Express Five medium (1 liter) was supplemented with 90 ml of L-Glutamine (200 mM, Invitrogen). During cultivation, cells were grown at 25 °C without CO2 in tissue culture flasks (75 cm2, Corning) and were split into fresh flasks when 90% confluent. The cells in passage 18 were seeded into 96-well plates (Falcon 96 Well Tissue Culture Plates, Corning) with 40,000 cells per well in 100 μl using an electronic multichannel pipette VIAFLO (1250 μ!, INTEGRA) on a pipetting robot ASSIST (INTEGRA). The cells 24 h growth rate and viability were monitored in the culture dishes (in duplicate; 100 mm, Corning) with 12×106 cells in 14 ml medium. Cell counting and assessment of cell viability were performed using the Cell Counter and Analyzer System (CASY, Roche).

#### Transient co-transfection

24 hours after cell plating, transient co-transfection on the robot system was performed using FuGENE® HD Transfection Reagent (Promega) according to the manufacturer’s protocol. To avoid multiple freeze-thaw processes, the renilla control plasmid and three inter-plate control plasmids (pUC19, pUG9, pZQ3) were aliquoted in PCR strips sufficient for one transfection experiment. The isolated reporter plasmids and inter-plate control plasmids were transferred into 96-well master mix plates according to the transfection plate layout together with renilla control plasmids (except for untreated cells (UTCs), reporter plasmid or inter-plate control plasmid renilla control plasmid ratio = 8: 1, total DNA amount 0.945 μg per well). Wells indicated with green shadows were filled with various reporter plasmids containing synthetic promoter constructs to be tested. 2.3 μl/well FuGENE® HD Transfection Reagent was added and the FuGENE® HD-DNA mixture was incubated for 5 min at RT (FuGENE® HD: DNA ratio ∼ 2.4: 1). 10 μl FuGENE® HD-DNA mixture was then added per well into 96-well cell culture plates. The transient co-transfections were performed in duplicates for cells with and without ecdysone treatment.

#### Ecdysone treatment

Cells were incubated for 22 h after transfection, followed by 2 h of ecdysone treatment (final ecdysone concentration: 10 μM; 20-Hydroxyecdysone, Sigma-Aldrich). The other replicate transfected cell plate was treated with the same volume (10 μl/well) of cell culture medium (Express Five medium supplemented with L-Glutamine) and incubated for 2 h.

#### Dual luciferase assay

40 μl/well of the mediums was removed from each cell culture plate and 20 μl/well of cells were transferred into the final readout plates. For each measurement, samples from four 96-well cell culture plates were joined into two 384-well plates (one for firefly luminescence measurement, the other for renilla luminescence measurement; AlphaPlate-384, PerkinElmer). ONE-GloTM Luciferase Assay System (Promega) and Renilla-Glo® Luciferase Assay System (Promega) were used respectively (reagent amount: 20 μl/well). There was a common crosstalk issue between two adjacent wells caused by the bleed-through of the stronger luminescence signal to the other. In the optimized protocol, firefly luminescence signal was measured twice with strong signals (> 2*105 RLU, relative light unit) identified in the first measurement and removed before the second measurement (samples were pipetted out and a highly-concentrated dye (1 mM Nile Blue A, Sigma-Aldrich) that quenches the luminescence signal was added instead). This experimental procedure was designed to solve the crosstalk issue between adjacent wells that we observed for strong promoters (Figure S1). Bioluminescence signals were measured using a SpectraMax Microplate Reader (Molecular Devices).

### Data analysis

#### Reads mapping for the sequencing results of the picked clones

Sequencing reads were demultiplexed based on the Illumina indexes and the designed barcodes in our customized primers. The most enriched sequence (at least 3-fold enrichment against the second most frequent sequence) for each sample was used and trimmed to match the target region of our synthetic promoter construct (part of *block 2,* blocks 3-6 and *block* 7). The trimmed reads were mapped to our designed library using the pairwise alignment method.

#### Data preprocessing and normalization

For each plate, firefly luciferase expression values (*FF*) of each tested samples were normalized to their renilla luciferase values (*REN*) as well as *FF* values of the inter-plate controls. The 1^st^ firefly measurements (FF1) were used as the readout values for samples with strong promoters (*FF1* > 2*10^5^ RLU) and the 2^nd^ firefly measurements (FF2, signal degradation corrected) were used for other weaker samples.

Background value (*BG*) was calculated as the arithmetic mean of negative control signals (pUC19 and UTCs) got from 2^nd^ firefly measurements (avoiding the potential crosstalk issue; Equation 1). Normalized value of positive control pUG9 (*Norm_pUG_*_9_) was defined as the arithmetic mean of its *FF1* signals with *BG* subtracted divided by its *REN* signals (Equation 2).

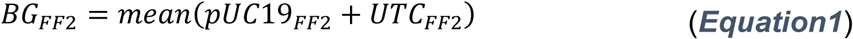

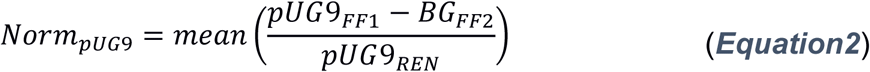

The final normalized luciferase expression value for each tested sample (*x*_i_) was calculated as Equation 3: its *FF*_*i*_ signal (*FF1* for strong promoters and *FF2* for others) with *BG* subtracted was firstly normalized to its *REN_i_* signal and then to the normalized control *Norm_PUG9_*; the value was then log_2_-transformed. This value was used as the estimate of the corresponding synthetic promoter activity.

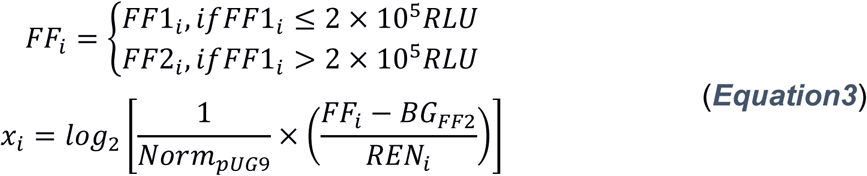

#### Outlier identification and filtering

We firstly filtered out samples with outlier renilla signals that we found out to be too high or too low to provide an accurate data normalization (*REN* > 10000 RLU or *REN* < 300 RLU, respectively), and then calculated the median and standard deviation (SD) for normalized luciferase signals of each promoter construct x (> 88% with at least three replicates for both with and without ecdysone stimulation). The score used for defining outliers was calculated as:

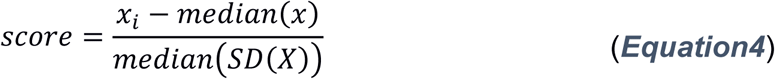

Here, *x,* as described above, represented the normalized expression value of /^1h^ replicate for construct x. *SD(X*) denoted all SDs for entire synthetic promoter construct library *X.* The scores with an absolute value of no less than 3 were labeled as outliers and were excluded from further analysis.

