## Supplemental figures and tables for "Large-scale analysis of *Drosophila* core promoter function using synthetic promoters"

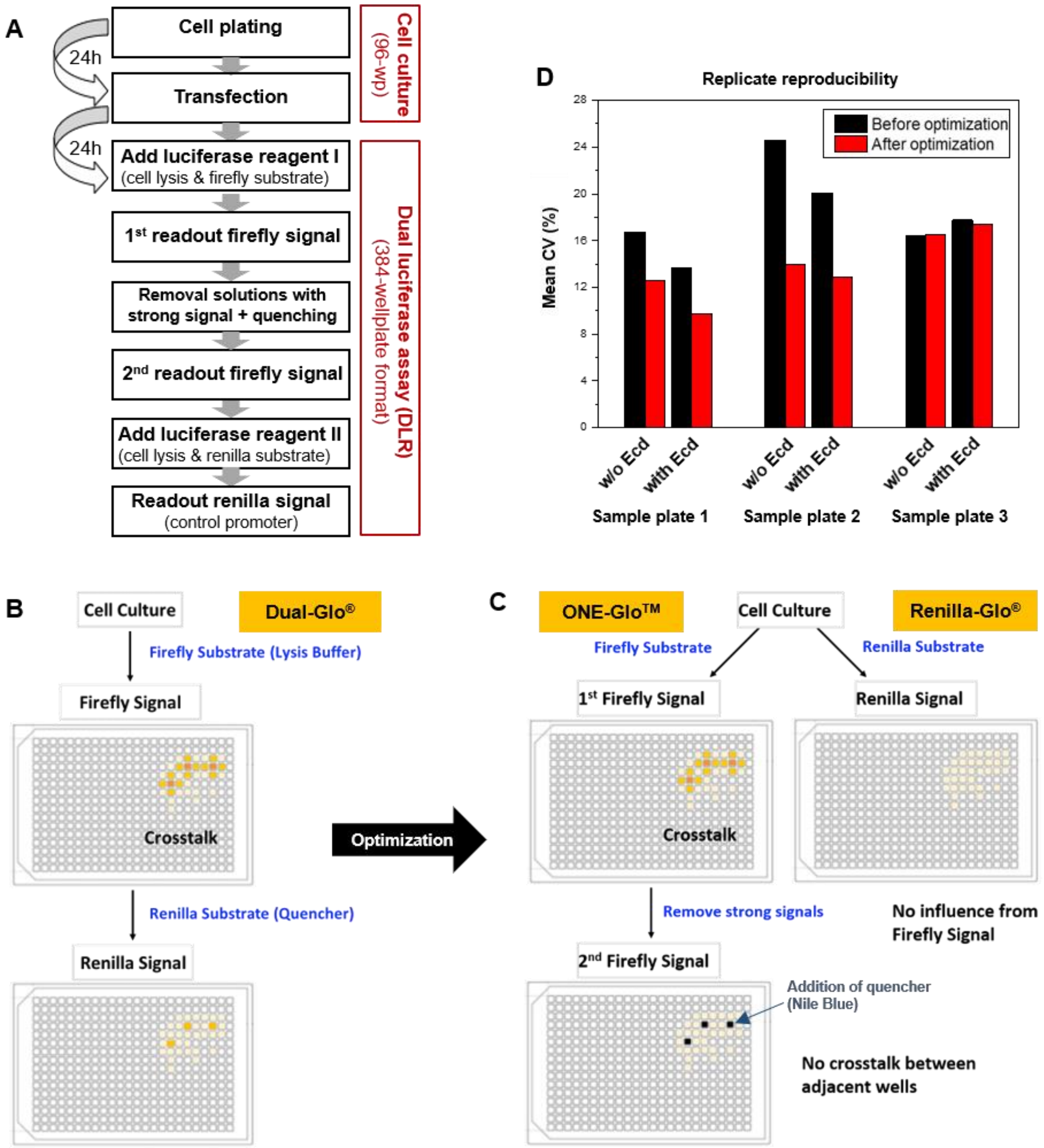

**Figure S1**. Assay development and reproducibility.

(A) Workflow for automated transfection in 96-wellplate format, followed by dual luciferase (DLR) assay in 384-wellplate format (**MATERIAL AND METHODS**). The transfection, lysis and cell detachement occurs in four cell culture 96-wellplates, followed by their splitting into two 384-wellplates for separate readout of the firefly and renilla luminescence signals. This enabled to gain 4-fold higher throughput and to save 2/3 luciferase assay reagent.

(B-C) Experimental strategy to eliminate *crosstalk* artefacts. Separating firefly and renilla readouts (the two upper panels) avoids potential *crosstalk* between the firefly and renilla luminescence light within the same well. A second readout of the firefly signal after removal of the solution from wells with very strong signal eliminates the *crosstalk* between neighboring wells.

(D) Comparison of expression measurements for three 96-well plates containing the same promoter construct samples measured on different days with and without ecdysone induction. Standard normalization (before optimization) uses only the ratios of firefly and renilla signals. After optimization of data normalization procedure of the luciferase assay readout (**MATERIAL AND METHODS**) the mean reproducibility of the measurements improved from a mean coefficient of variation ~13% after optimizations, versus ~18% before.

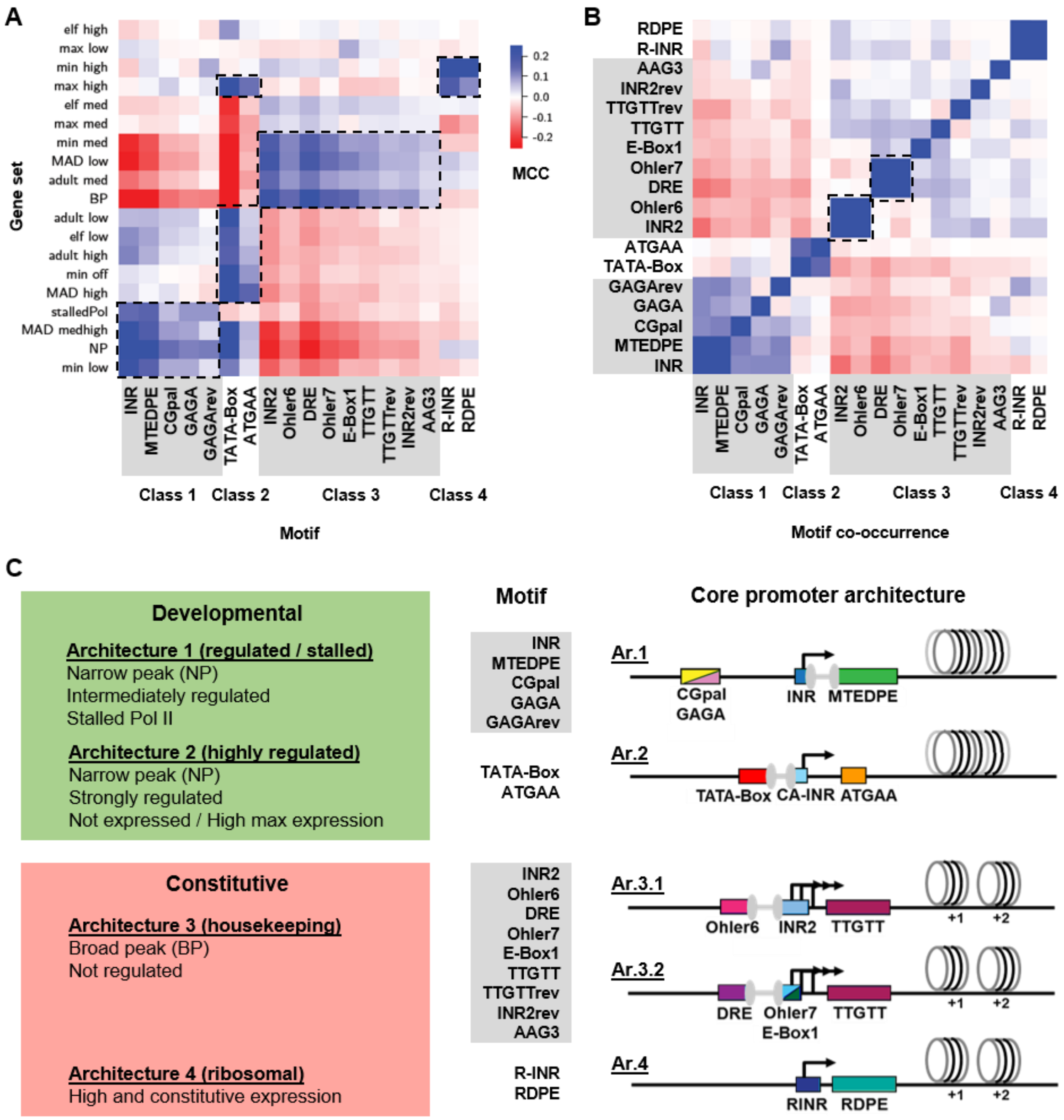

A
elf high
max low
min high
max high
elf med
max med
min med
MAD low
adult med
BP
adult low
elf low
adult high
min off
MAD high
stalledPol
MAD medhigh
NP
min low
Gene set
INR
MTEDPE
CGpal
GAGA
GAGArev
TATA-Box
ATGAA
INR2
Ohler6
DRE
Ohler7
E-Box1
TTGTT
TTGTTrev
INR2rev
AAG3
R-INR
RDPE
Class 1
Class 2
Class 3
Class 4
Motif
0.2
0.1
0.0
-0.1
-0.2
MCC
B
RDPE
R-INR
AAG3
INR2rev
TTGTTrev
TTGTT
E-Box1
Ohler7
DRE
Ohler6
INR2
ATGAA
TATA-Box
GAGArev
GAGA
CGpal
MTEDPE
INR
Class 1
Class 2
Class 3
Class 4
Motif co-occurrence
C
Developmental
Architecture 1 (regulated / stalled)
Narrow peak (NP)
Intermediately regulated
Stalled Pol II
Architecture 2 (highly regulated)
Narrow peak (NP)
Strongly regulated
Not expressed / High max expression
Constitutive
Architecture 3 (housekeeping)
Broad peak (BP)
Not regulated
Architecture 4 (ribosomal)
High and constitutive expression
Motif
INR
MTEDPE
CGpal
GAGA
GAGArev
TATA-Box
ATGAA
INR2
Ohler6
DRE
Ohler7
E-Box1
TTGTT
TTGTTrev
INR2rev
AAG3
R-INR
RDPE
Core promoter architecture
Ar.1
CGpal
GAGA
INR
MTEDPE
Ar.2
TATA-Box
CA-INR
ATGAA
Ar.3.1
Ohler6
INR2
TTGTT
+1
+2
Ar.3.2
DRE
Ohler7
E-Box1
TTGTT
+1
+2
Ar.4
RINR
RDPE
+1
+2

**Figure S2**. Genome-wide Analysis of Promoter Features.

(A) Correlation of core promoter motifs to 19 gene sets with different features (motifs and meaning of the different features listed in **Table S1**) reveals four distinct motif classes: class 1 motifs enriched in the gene sets of stalledPol, MAD medhigh, NP and min low; class 2 motifs enriched in the gene sets of max high, adult low, elf low, adult high, min off and MAD high; class 3 motifs enriched in the gene sets of min med, MAD low, adult med and BP; class 4 motifs enriched in the gene sets of min high and max high (details on the motifs in **Table S2** and on the different gene sets in **Table S3**). MCC: Matthews correlation coefficient. Class 1 motifs (INR, MTEDPE, CGpal, GAGA, GAGArev) occur in genes with NP core promoters. The enriched genes are intermediately regulated and show strong correlations to stalled Pol II. Class 2 motifs including TATA-Box and ATGAA also present in NP promoter genes, however, the enriched genes are strongly regulated ones that are either not expressed or most highly expressed in at least one developmental stage. Class 3 motifs (INR2, Ohler6, DRE, Ohler7, E-Box1, TTGTT, TTGTTrev, INR2rev, AAG3) are the ones only found in genes with BP core promoters. The enriched genes are not regulated and similarly expressed in all developmental stages (housekeeping function). Class 4 motifs (R-INR, RDPE) correlate with strongly expressed genes which mainly encode the ribosomal proteins.

(B) Correlation of core promoter motifs to each other indicates motif co-occurrence within the same promoter, which agrees with the four defined classes and also suggests two distinct preferred compositions in the 3rd class (INR2 + Ohler6 pair and DRE + Ohler7 pair).

(C) Based on the motif classes identified by *XXmotif*, four core promoter architectures (Ar. 1 – Ar. 4) are defined accordingly, reflecting different modes of transcriptional regulation at the core promoter. They are named as regulated/stalled, highly regulated, housekeeping and ribosomal architectures. They can be further grouped into developmental (Ar.1, Ar.2; highlighted in green) and constitutive (Ar.3.1, Ar.3.2, Ar.4; highlighted in red) core promoters based on their association with gene functions. The motif co-occurrence analyzed in (A-B) agrees with the four defined classes and also suggests two distinct preferred compositions in the 3rd class (INR2 + Ohler6 pair and DRE + Ohler7 pair, see (B).

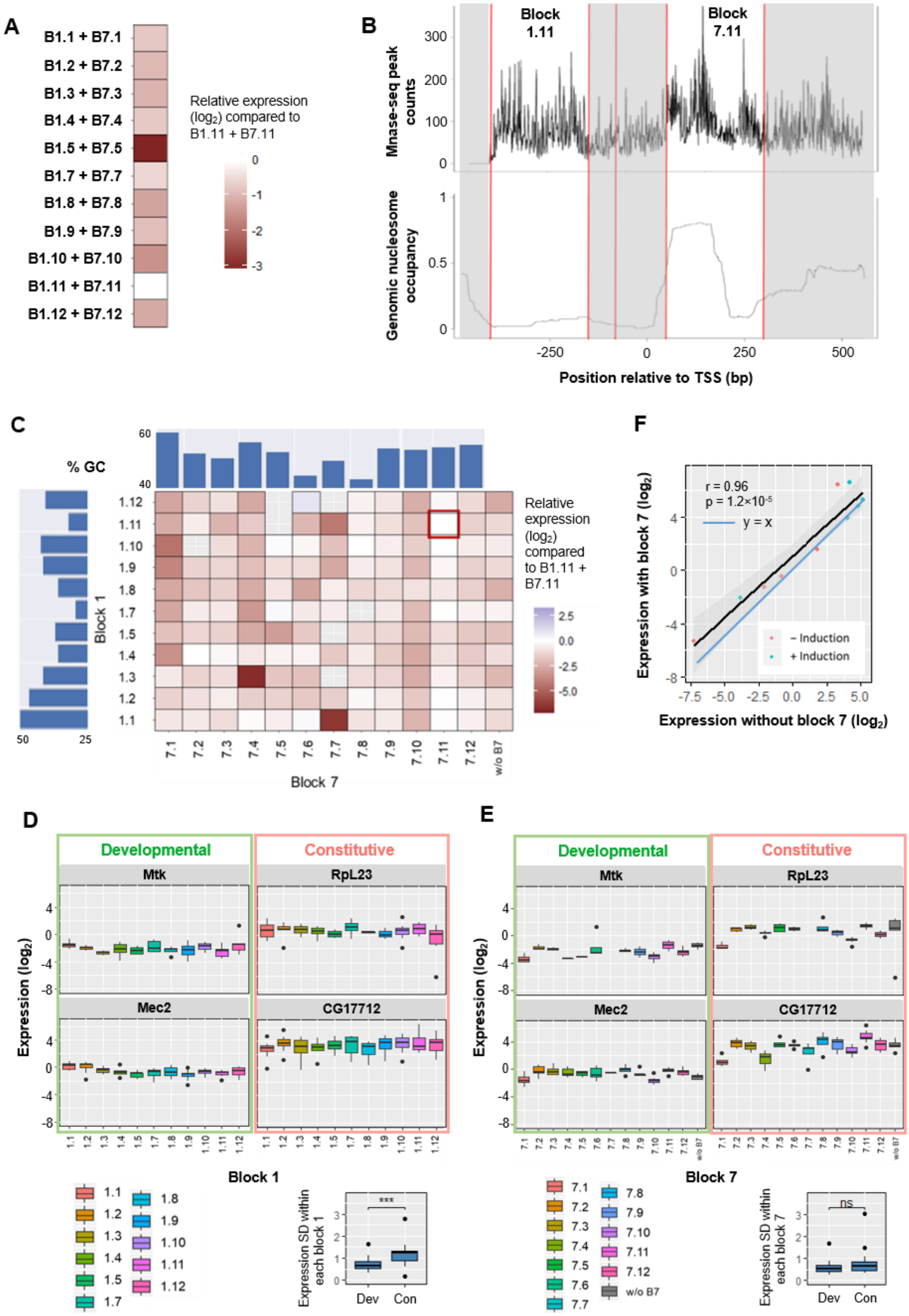
A
B1.1 + B7.1
B1.2 + B7.2
B1.3 + B7.3
B1.4 + B7.4
B1.5 + B7.5
B1.7 + B7.7
B1.8 + B7.8
B1.9 + B7.9
B1.10 + B7.10
B1.11 + B7.11
B1.12 + B7.12
Relative expression (log2) compared to B1.11 + B7.11
0
-1
-2
-3
B
Block 1.11
Block 7.11
Mnase-seq peak counts
300
200
100
0
Genomic nucleosome occupancy
1
0.5
0
-250
0
250
500
Position relative to TSS (bp)
C
% GC
60
40
1.12
1.11
1.10
1.9
1.8
1.7
1.5
1.4
1.3
1.2
1.1
Block 1
50
25
7.1
7.2
7.3
7.4
7.5
7.6
7.7
7.8
7.9
7.10
7.11
7.12
w/o B7
Block 7
Relative expression (log2) compared to B1.11 + B7.11
2.5
0.0
-2.5
-5.0
F
r = 0.96
p = 1.2×10-5
y = x
– Induction
+ Induction
Expression with block 7 (log2)
Expression without block 7 (log2)
-7.5
-5.0
-2.5
0.0
2.5
5.0
D
Developmental
Constitutive
Mtk
RpL23
Mec2
CG17712
Expression (log2)
Block 1
1.1
1.2
1.3
1.4
1.5
1.7
1.8
1.9
1.10
1.11
1.12
Expression SD within each block 1
***
Dev
Con
E
Developmental
Constitutive
Mtk
RpL23
Mec2
CG17712
Expression (log2)
Block 7
7.1
7.2
7.3
7.4
7.5
7.6
7.7
7.8
7.9
7.10
7.11
7.12
w/o B7
Expression SD within each block 7
ns
Dev
Con

**Figure S3**. The effect of nucleosomal context on expression.

(A) Heatmap depicting the relative expression measurements of promoter constructs with different pairs of block 1 and block 7 compared to *B1.11* + *B7.11* expressions ($log_2$ scale). Results were pooled for all tested native core promoters to calculate the average deviation to *B1.11* + *B7.11* expressions.

(B) Upper panel: MNase-seq measurement of the nucleosome occupancy for one synthetic promoter construct (*block 1.11* + *block 2* + *MED4 (native)* + *block 7.11*). The peak in *block 7.11* (highlighted in white on the right) suggests the presence of a +1 nucleosome; lower panel: nucleosome occupancy for sequences at similar genomic locations of the gene CG8613, from which *block 1.11* and *block 7.11* sequences were derived

(C) Heatmap depicting the relative expression measurements of promoter constructs with different free combinations of *block 1* and *block 7* compared to *B1.11* + *B7.11* expressions (marked with a red rectangle). Results were pooled for all tested native core promoters to calculate the average deviation to *B1.11* + *B7.11* expressions. Bar plots on the top and the left represent the GC content of each *block 1* and *block 7* sequence. *Block 7* with column "w/o B7" represents the results obtained from promoters without block 7 sequence.

(D) Boxplots depicting *block 1* effects for tested core promoters. Effects of different *block 7*s were merged in each column (within the same *block 1*): the median SD is 0.66 for developmental promoters compared to 1.23 for constitutive promoters (lower right corner); Wilcoxon rank-sum test p = $3.1\times10^{-4}$, significant.

(E) Boxplots depicting *block 7* effects for tested core promoters. Effects of different *block 1*s were merged in each column (within the same *block 7*): the median SD is 0.54 for developmental promoters compared to 0.64 for constitutive promoters (lower right corner); Wilcoxon rank-sum test p = 0.3, not significant. Block 7 with column "w/o B7" represents the results obtained from promoters without block 7 sequence. Developmental and constitutive promoters are highlighted in green and red, respectively.

(F) Comparison of the expression measurements for promoter variants with *block 7.11* versus without *block 7*. Expressions with and without ecdysone induction are labeled cyan and red, respectively. Black line: linear regression (with 95% confidence interval shown in gray, PCC r = 0.96, p = $1.2\times10^{-5}$). Blue line: y = x.

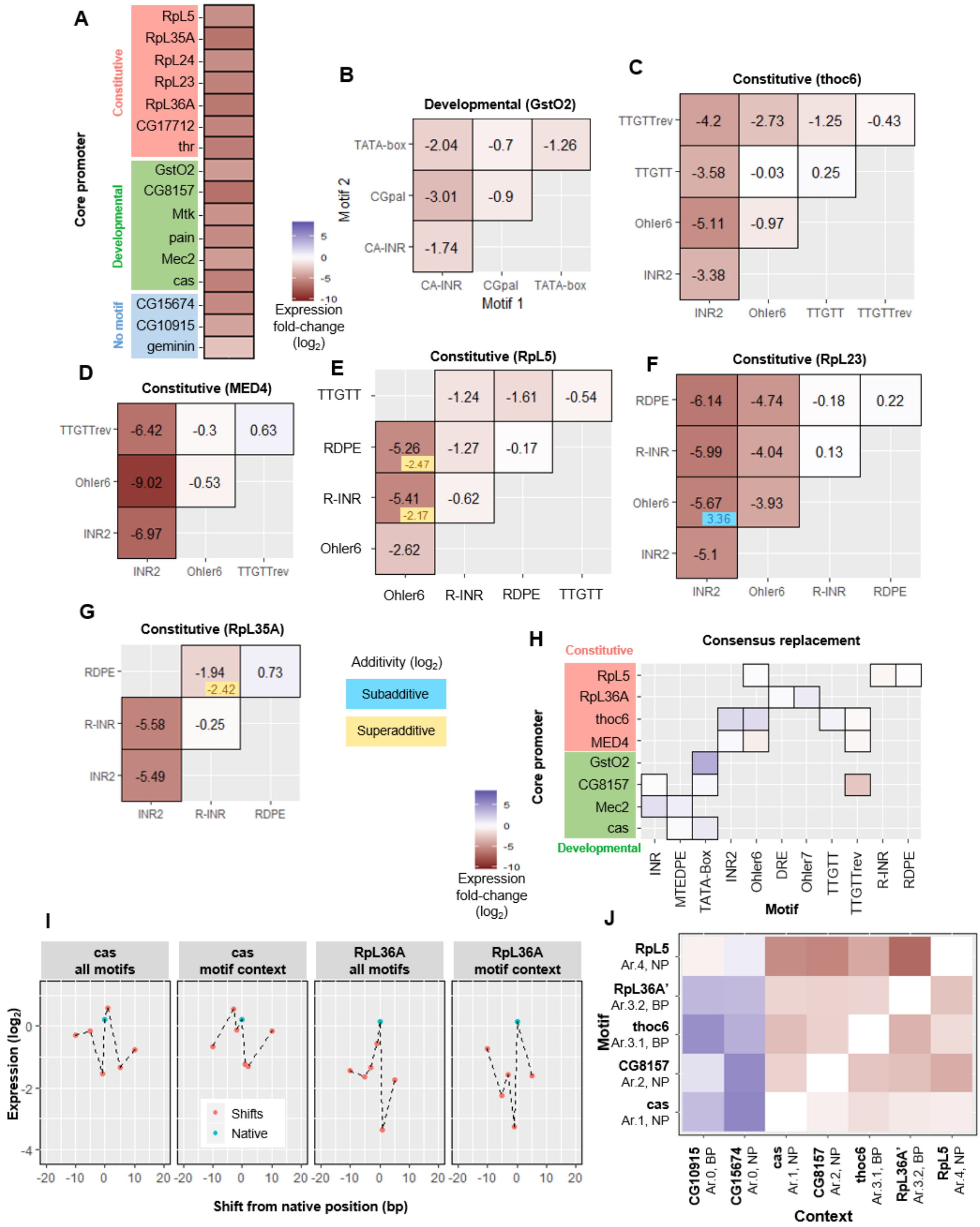
A
Core promoter
Constitutive
RpL5
RpL35A
RpL24
RpL23
RpL36A
CG17712
thr
Developmental
GstO2
CG8157
Mtk
pain
Mec2
cas
No motif
CG15674
CG10915
geminin
Expression fold-change (log2)
B
Developmental (GstO2)
Motif 2
TATA-box
CGpal
CA-INR
-2.04
-0.7
-1.26
-3.01
-0.9
-1.74
CA-INR
CGpal
TATA-box
Motif 1
C
Constitutive (thoc6)
TTGTTrev
TTGTT
Ohler6
INR2
-4.2
-2.73
-1.25
-0.43
-3.58
-0.03
0.25
-5.11
-0.97
-3.38
INR2
Ohler6
TTGTT
TTGTTrev
D
Constitutive (MED4)
TTGTTrev
Ohler6
INR2
-6.42
-0.3
0.63
-9.02
-0.53
-6.97
INR2
Ohler6
TTGTTrev
E
Constitutive (RpL5)
TTGTT
RDPE
R-INR
Ohler6
-1.24
-1.61
-0.54
-5.26
-2.47
-1.27
-0.17
-5.41
-2.17
-0.62
-2.62
Ohler6
R-INR
RDPE
TTGTT
F
Constitutive (RpL23)
RDPE
R-INR
Ohler6
INR2
-6.14
-4.74
-0.18
0.22
-5.99
-4.04
0.13
-5.67
3.36
-3.93
-5.1
INR2
Ohler6
R-INR
RDPE
G
Constitutive (RpL35A)
RDPE
R-INR
INR2
-1.94
-2.42
0.73
-5.58
-0.25
-5.49
INR2
R-INR
RDPE
Additivity (log2)
Subadditive
Superadditive
Expression fold-change (log2)
H
Consensus replacement
Constitutive
RpL5
RpL36A
thoc6
MED4
Developmental
GstO2
CG8157
Mec2
cas
Core promoter
INR
MTEDPE
TATA-Box
INR2
Ohler6
DRE
Ohler7
TTGTT
TTGTTrev
R-INR
RDPE
Motif
I
cas all motifs
cas motif context
RpL36A all motifs
RpL36A motif context
Expression (log2)
Shifts
Native
Shift from native position (bp)
J
Motif
RpL5 Ar.4, NP
RpL36A' Ar.3.2, BP
thoc6 Ar.3.1, BP
CG8157 Ar.2, NP
cas Ar.1, NP
CG10915 Ar.0, BP
CG15674 Ar.0, NP
cas Ar.1, NP
CG8157 Ar.2, NP
thoc6 Ar.3.1, BP
RpL36A' Ar.3.2, BP
RpL5 Ar.4, NP
Context

**Figure S4**. All-motifs and pairwise knockouts, consensus replacement, all-motifs shifts, and motif context exchange.

(A) Heatmap depicting the mean expression levels of promoter constructs with all-motif knockouts ($log_2$ scale) for constitutive and developmental promoters. The expression levels dramatically decrease for all investigated promoters. The expression levels of the motifless promoters are indicated on the bottom for comparison. Constitutive, developmental and motifless promoters are highlighted in red, green and blue, respectively.

(B-G) The effect of pairwise motif knockout in different core promoters ($log_2$ scale). Heatmaps of the mean expression fold changes compared to wild-type expressions for pairwise knockout of motifs compared to individual knockouts (diagonals) in GstO2 (B), thoc6 (C), MED4 (D), RpL5 (E), RpL23 (F), and RpL35A (G), respectively. Additivity was calculated as the difference between the pairwise effect and the sum of two individual effects. Subadditive (blue): > 0; superadditive (yellow): < 0; effects > 3×SD shown in the corner of each pairwise effect.

(H) Consensus replacement. Heatmap depicting the mean expression fold changes compared to wild-type expressions after replacing with motif consensus sequences derived by XXmotif. Constitutive and developmental promoters are highlighted in red and green, respectively.

(I) Effect of all-motif shifts and context sequence shifts. The expression measurements of natives (cyan dots) and positional shifts (red dots) of either all motifs or motif context sequence in cas and RpL36A, the two core promoter configurations also used for testing the individual motif shifts.

(J) Effect of motif context sequence exchange. Heatmap depicting the mean expression fold changes caused by motifs (y-axis) inserting of RpL5, RpL36A, thoc6, CG8157, and cas to different context sequences (x-axis). The heatmap shows the expression changes relative to wild-type expressions of the context-origin promoters CG10915, CG15674, cas, CG8157, thoc6, RpL36A’, and RpL5, respectively.

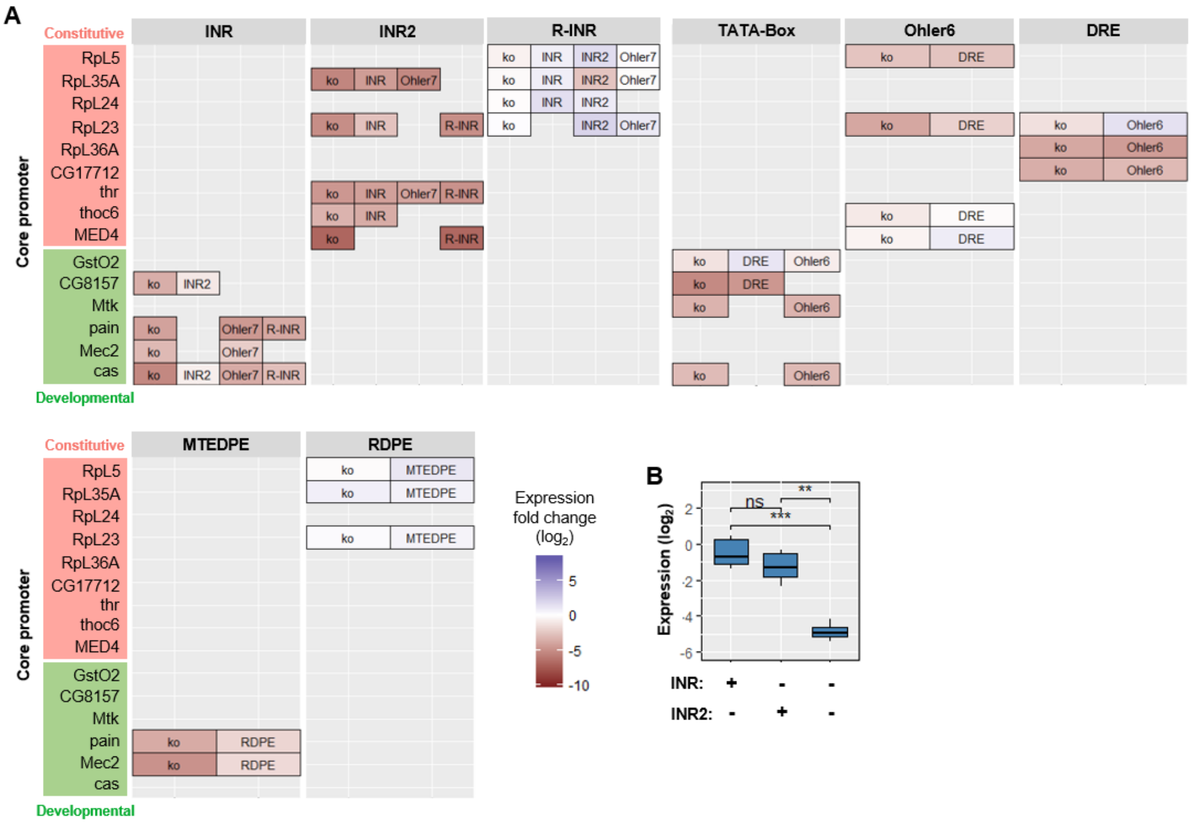

**Figure S5**. The effect of motif substitutions.

(A) Heatmap depicting the mean expression fold changes compared to wild-type expressions for motif knockout and substitution with positionally or functionally equivalent motifs from other architectures. Constitutive and developmental promoters are highlighted in red and green, respectively.

(B) Boxplot depicting the effects of INR being substituted by INR2 in cas and CG8157 (all measurements in these two core promoter constructs were pooled together; Wilcoxon rank-sum test p = 0.0051 for comparing substitution with knockout (significant) and p = 0.17 for comparing substitution with wild-type (not significant).

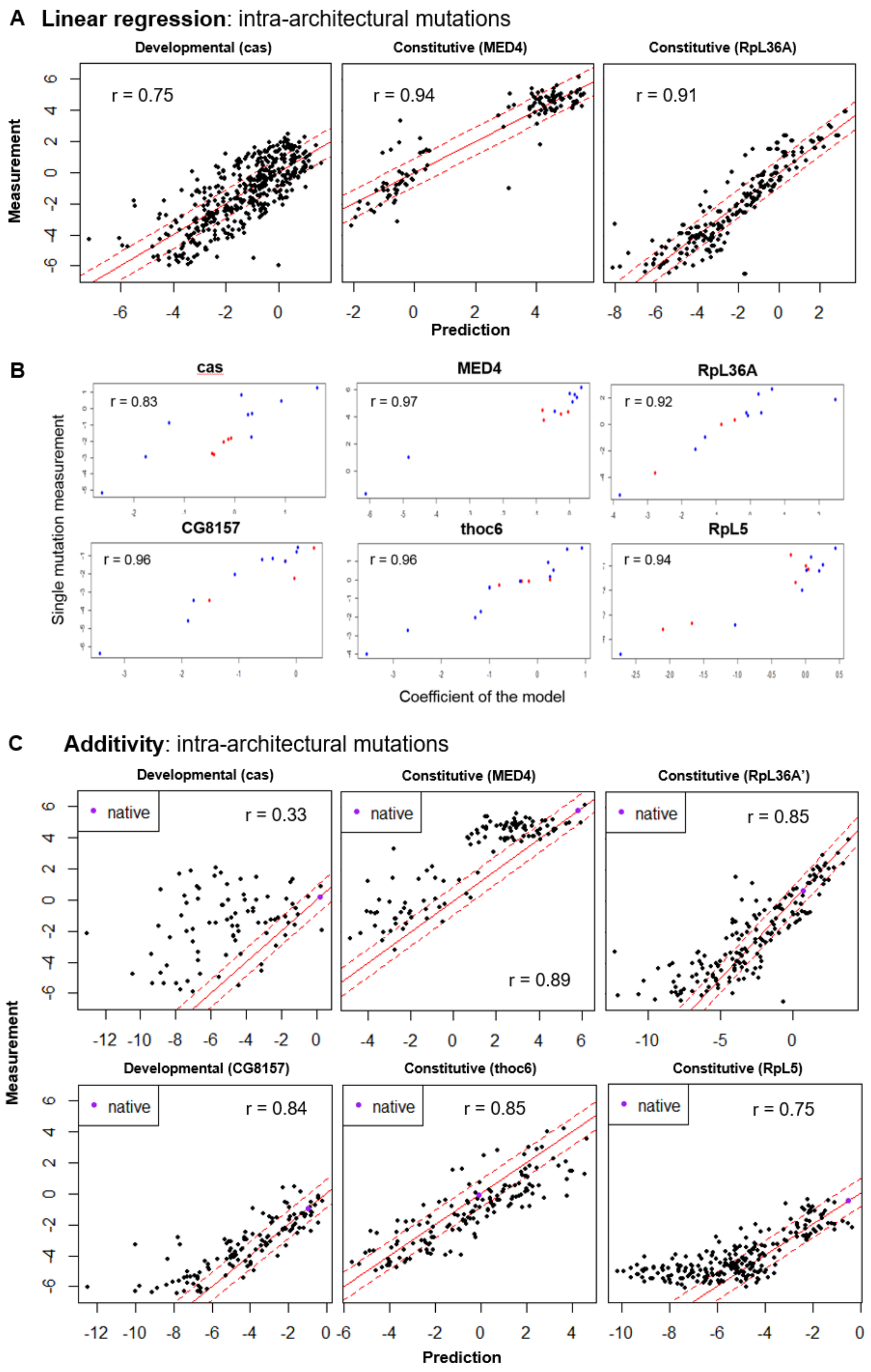
A Linear regression: intra-architectural mutations
Developmental (cas)
Constitutive (MED4)
Constitutive (RpL36A)
r = 0.75
r = 0.94
r = 0.91
Measurement
Prediction
B
cas
MED4
RpL36A
r = 0.83
r = 0.97
r = 0.92
CG8157
thoc6
RpL5
r = 0.96
r = 0.96
r = 0.94
Single mutation measurement
Coefficient of the model
C Additivity: intra-architectural mutations
Developmental (cas)
Constitutive (MED4)
Constitutive (RpL36A')
native
r = 0.33
r = 0.89
r = 0.85
Developmental (CG8157)
Constitutive (thoc6)
Constitutive (RpL5)
r = 0.84
r = 0.85
r = 0.75
Measurement
Prediction

**Figure S6**. Linear regression and additive models applied to predict the synthetic promoter activity based on individual motif features.

(A) Linear regression applied to the intra-architectural mutations for cas, MED4 and RpL36A. The measured expressions (on the y-axis) of the promoters with combinatorial motif mutations compared to the predicted expressions (on the x-axis) from the linear regression ($log_2$ scale). Red solid line: y = x; red dashed lines: $y = x \pm 3 \times SD_{noise}$ ($SD_{noise}$ denoted the experimental noise, that is the median of all SDs for all measured synthetic promoter constructs).

(B) Additive models applied to the same intra-architectural mutations for the 6 tested core promoters to predict synthetic promoter activity based on individual measured motif features. The measured expressions (on the y-axis) sequences with combinatorial motif mutations compared to the predicted expressions (on the x-axis) from the additive model ($log_2$ scale), which consists of adding directly the measured expressions of the individual features. Red solid line: y = x; red dashed lines: $y = x \pm 3 \times SD_{noise}$. Purple dot: the native expression of each promoter.

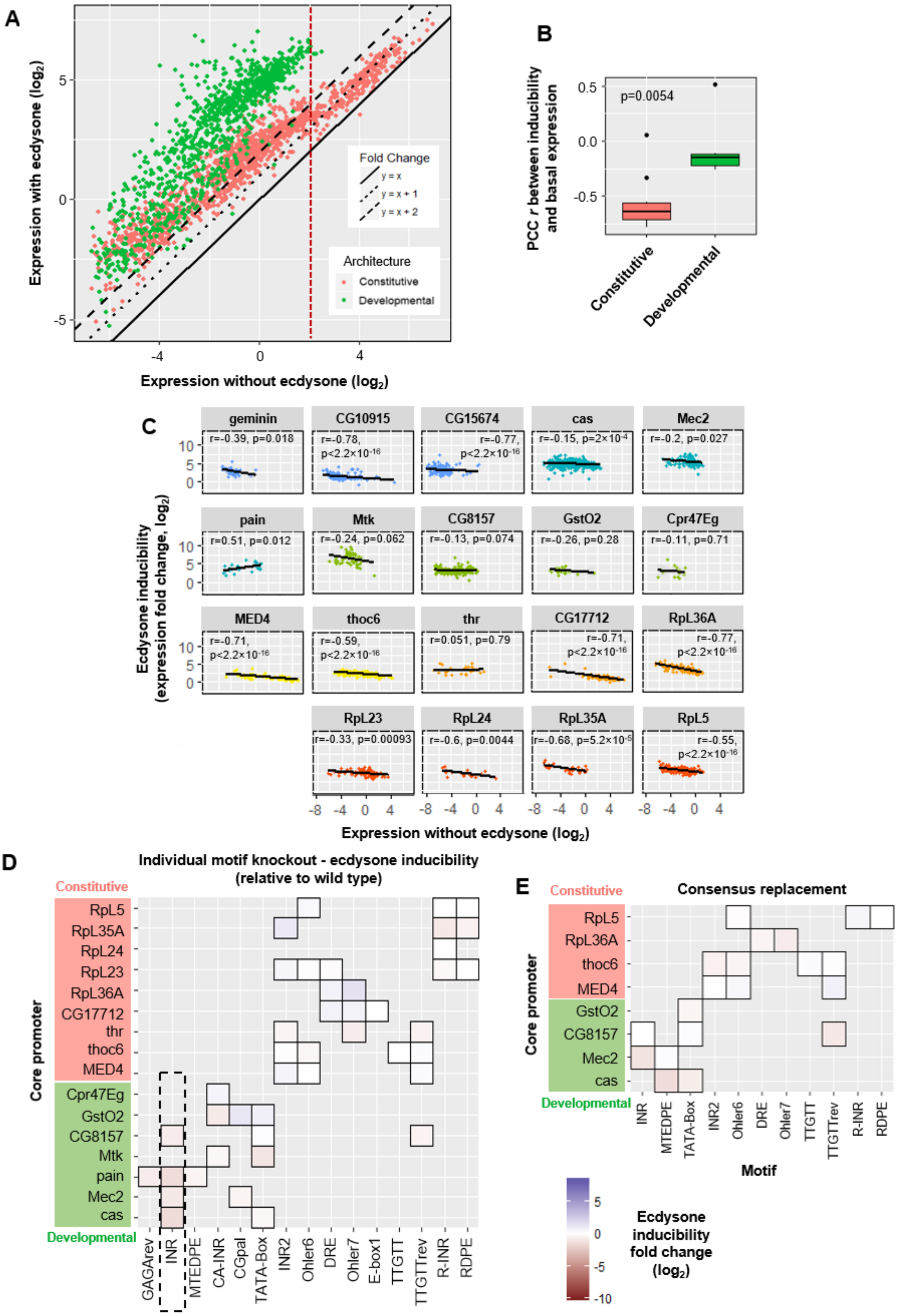
A
Expression with ecdysone (log2)
Expression without ecdysone (log2)
Fold Change
y = x
y = x + 1
y = x + 2
Architecture
Constitutive
Developmental
B
PCC r between inducibility and basal expression
p=0.0054
Constitutive
Developmental
C
Ecdysone inducibility (expression fold change, log2)
Expression without ecdysone (log2)
geminin
r=-0.39, p=0.018
CG10915
r=-0.78, p<2.2×10-16
CG15674
r=-0.77, p<2.2×10-16
cas
r=-0.15, p=2×10-4
Mec2
r=-0.2, p=0.027
pain
r=0.51, p=0.012
Mtk
r=-0.24, p=0.062
CG8157
r=-0.13, p=0.074
GstO2
r=-0.26, p=0.28
Cpr47Eg
r=-0.11, p=0.71
MED4
r=-0.71, p<2.2×10-16
thoc6
r=-0.59, p<2.2×10-16
thr
r=0.051, p=0.79
CG17712
r=-0.71, p<2.2×10-16
RpL36A
r=-0.77, p<2.2×10-16
RpL23
r=-0.33, p=0.00093
RpL24
r=-0.6, p=0.0044
RpL35A
r=-0.68, p=5.2×10-5
RpL5
r=-0.55, p<2.2×10-16
D
Individual motif knockout - ecdysone inducibility (relative to wild type)
Constitutive
Developmental
Core promoter
RpL5
RpL35A
RpL24
RpL23
RpL36A
CG17712
thr
thoc6
MED4
Cpr47Eg
GstO2
CG8157
Mtk
pain
Mec2
cas
GAGArev
INR
MTEDPE
CA-INR
CGpal
TATA-Box
INR2
Ohler6
DRE
Ohler7
E-box1
TTGTT
TTGTTrev
R-INR
RDPE
Motif
E
Consensus replacement
Constitutive
Developmental
Core promoter
RpL5
RpL36A
thoc6
MED4
GstO2
CG8157
Mec2
cas
INR
MTEDPE
TATA-Box
INR2
Ohler6
DRE
Ohler7
TTGTT
TTGTTrev
R-INR
RDPE
Motif
Ecdysone inducibility fold change (log2)

**Figure S7**. Ecdysone induction effect ($log_2$ scale) grouped by promoter architectures.

(A) Scatterplot depicting the expression measurements with ecdysone induction versus measurements without ecdysone for all tested promoters separated by core promoter architectures. Constitutive and developmental promoters are plotted in red and green, respectively. Three types of line are used to indicate the expression fold change with no increase ($y = x$; solid line), 2-fold increase ($y = x + 1$; dotted line) and 4-fold increase ($y = x + 2$; dashed line).

(B) Comparison of the PCCRs in (A) grouped by constitutive and developmental core promoters. Wilcoxon rank-sum test $p = 0.0054$.

(C) Expression fold changes (ecdysone inducibility) versus measurements without ecdysone and grouped by native core promoter sequences. The colors refer to different core promoter architectures. Black line: linear regression (with the 95% confidence interval shown in gray, PCC r and p are shown for each group).

(D) Heatmap depicting the ecdysone inducibility fold changes caused by individual knockout of motifs in different core promoters. Disrupted INR (highlighted with the black dotted line rectangle) had a slightly negative effect on changing the core promoter responsiveness to ecdysone. (~ 2.3-fold reduction on average, Wilcoxon rank-sum test $p = 2.1\times10^{-5}$). Constitutive and developmental promoters highlighted in red and green, respectively.

(E) Heatmap depicting the ecdysone inducibility fold changes caused by consensus replacement of motifs in different core promoters. Constitutive and developmental promoters highlighted in red and green, respectively.

### Supplemental Tables

| | Motif Name | Sequence Logo | Distribution | Start Range | Gene Set |
|---|---|---|---|---|---|
| Known Motifs | INR* | 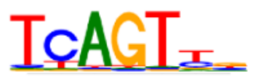 | 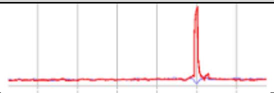 | -2 … -1 | NP |
| | MTE/DPE* | 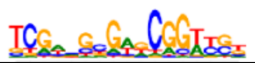 | 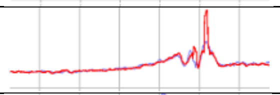 | 16 … 18 | NP |
| | GAGA* | 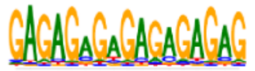 | 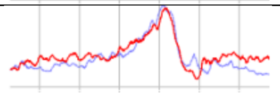 | -100 … -33 | NP |
| | GAGArev* | 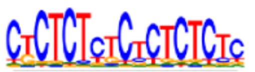 | 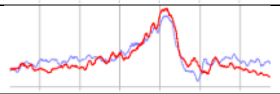 | -100 … 1 | NP |
| | INR2* | 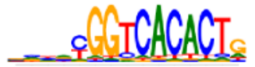 | 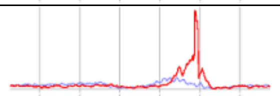 | -60 … 20 | BP |
| | DRE* | 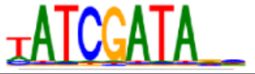 | 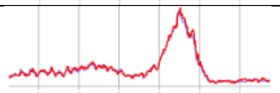 | -100 … -7 | BP |
| | Ohler7* | 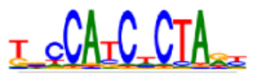 | 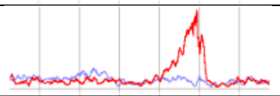 | -72 … 14 | BP |
| | E-Box1* | 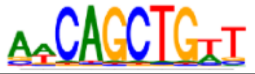 | 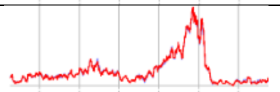 | -59 … 32 | BP |
| | Ohler6* | 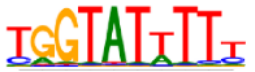 | 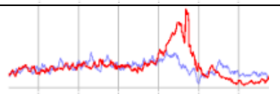 | -100 … -10 | BP |
| | TATA-Box* | 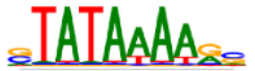 | 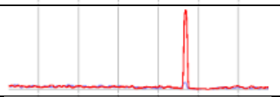 | -35 … -29 | MAD high |
| | R-INR* | 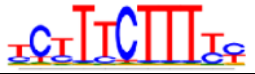 | 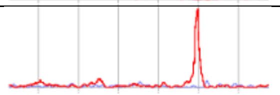 | -5 … -5 | min high |
| | E-Box2 | 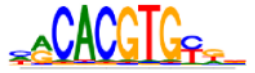 | 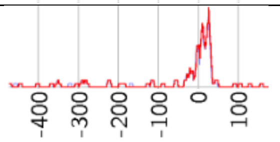  | -22 … 40 | elf high |
| Newly discovered | CGpal* | 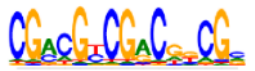 | 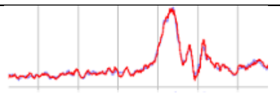 | -100 … -20 | NP |
| | INR2rev | 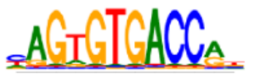 | 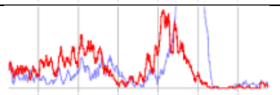 | -100 … -2 | BP |
| | TTGTT* | 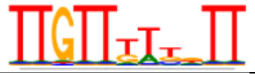 | 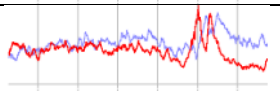 | -32 … 40 | MAD low |
| | TTGTTrev* | 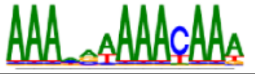 | 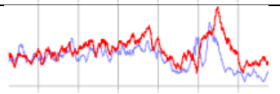 | -14 … 38 | BP |
| | AAG3 | 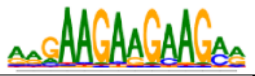 | 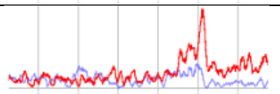 | -57 … 38 | min med |
| | ATGAA | 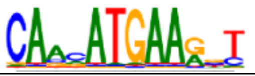 | 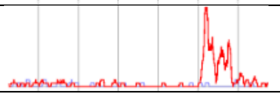 | -5 … 39 | MAD high |

| | | | | | |
|---|---|---|---|---|---|
| | RDPE* | 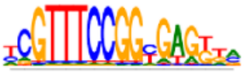 | 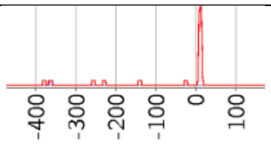 | 8 … 12 | min high |

| Gene Set | Definition |
|---|---|
| NP | TSS cluster width: narrow peak |
| BP | TSS cluster width: broad peak |
| MAD low | Mean absolute deviation (MAD) expression: low |
| MAD medhigh | MAD expression: medium to high (top 40% of genes) |
| MAD high | MAD expression: high (top 10% of genes) |
| min off | Minimum gene expression: nearly no expression |
| min low | Minimum gene expression: low |
| min med | Minimum gene expression: medium |
| min high | Minimum gene expression: high |
| max low | Maximum gene expression: low |
| max med | Maximum gene expression: medium |
| max high | Maximum gene expression: high |
| elf low | Gene expression in embryo, larva or female: low |
| elf med | Gene expression in embryo, larva or female: medium |
| elf high | Gene expression in embryo, larva or female: high |
| adult low | Gene expression in adult: low |
| adult med | Gene expression in adult: medium |
| adult high | Gene expression in adult: high |
| stalledPol | All genes classified as stalled by (Hendrix et al., 2008) |

**Table S1.** Core promoter motifs detected by *XXmotif* in *D.mel*.

Upper panel: known motifs in literature (on the top), and newly discovered motif candidates (on the bottom). Motifs highlighted with "*" are the ones used in this study. "Distribution" depicts the distribution of all assigned motifs within the most enriched gene set (details in **Table S2**) smoothed over five nucleotides. "Start Range" is the region that the motif's 1st nucleotide most locates relative to the defined TSS.

Lower panel: Legend for the abbreviations used in the column "Gene Set".

| Motif Name | Start Position | End Position | Max Score | Threshold Score |
|---|---|---|---|---|
| CA-INR | -3 | -3 | 16.28 | 6.5 |
| CGpal | -100 | -20 | 26.46 | 3.7 |
| DRE | -100 | -7 | 15.78 | 7.3 |
| E-Box1 | -59 | 32 | 16.66 | 11.6 |
| GAGA | -100 | -33 | 31.27 | 0 |
| GAGArev | -100 | 1 | 29.40 | 2 |
| INR | -2 | -1 | 11.15 | 4.6 |
| INR2 | -60 | 20 | 22.42 | 8.1 |
| MTEDPE | 16 | 18 | 24.03 | 0.6 |
| Ohler6 | -100 | -10 | 15.94 | 7.5 |
| Ohler7 | -72 | 14 | 19.30 | 7.3 |
| RDPE | 8 | 12 | 28.38 | 14 |
| R-INR | -5 | -5 | 16.77 | 5 |
| TATA-Box | -35 | -29 | 14.73 | 5.5 |
| TTGTT | -32 | 40 | 15.94 | 1.6 |
| TTGTTrev | -14 | 38 | 18.56 | 2.2 |

**Table S2.** A summary of *XXmotif*-annotated core promoter motif features used in this work**.**

Motif 1$^{st}$ nucleotide locates within the range between “Start Position” and “End Position” relative to the defined TSS (also shown in the column “Start Range” in **Table S1**). “Max Score” is the PWM score of the motif consensus. “Threshold Score” is the minimal score that maximizes the mutual information between a certain motif and all positively correlated gene sets.

| Native Promoter | Sequence (5’-3’) | TSS Distribution |
|---|---|---|
| FBgn0003701 | TGTAGTATTTGTGTTCTGATATGAAAACTAGAGATATCGATGTTATCGTTA<br>AAGGATTTCCAGCTTTAGCATGGACGGTCACACTGGATCTCAAAATCTG<br>GCGCAAAGCAACAAAAAAGGAAGCGTCGCAG | BP |
| FBgn0004878 | CGGTGGCGAGAGGGTTGCACTTGGGCATGGATTTGCGCAATTTTTGCTA<br>TATTAGCCGGAGGCCGCGGAAGAGTTGAAGCAGTTTGAGCCTCGCAGC<br>CGAACTTTGAGGATCGCTGAGACGAGACGCCGTG | NP |
| FBgn0010078 | CTTGGTTATAATTAGGTTATTTTTTCGATATTTTGAGGTATATTTCTACGAT<br>AGATCGGCGGTCACATCGTATTTCCCTCCTTTTCGTTTTCGTTTCCGGCG<br>AAGTAAGTATAATAAAATCTCCACGTTTT | NP |
| FBgn0014865 | CTCAAATAAAAAGTCCCCAATCTGCGACTCGTTTGTCTGGGACTGAGCTA<br>TAAAAGCCTCACCATCTCAACGCTCAAAGCATCAATCAATTCCCGCCACC<br>GAGCTAAGtaGCAACTTAATCTTGGAGCGAT | NP |
| FBgn0027597 | TTTAAATAGATTTAGCTAGAAAATAGCTGACAGACACATATCGATATATCG<br>CTGCGATAGCCACAGCTGTTCACGCCCGCAGTTTAAGCGtaGTGGCAGC<br>CCTGGTCGGCCACCAAAAAATAAACATTGGA | BP |
| FBgn0030993 | GAGAGAACCAGTGCGCTCTTATCACGTGAGAACGCTTTTGGGCATTCAG<br>TTTGGCTTTTGCGGCGCTGACCGCTGGCGACAGTTTCGAATCCATAGCC<br>GATCGGAGAGCAACGAACGTAGGCCAGAACGGA | NP |
| FBgn0031980 | CATATCAAGTCACAACAATGAAACGAAAACCTATCGATAGCGCATGGCTT<br>GACGGCACGCTGCCATCGCTATGTGATTTCCTTCTTTTTTCGCCTTCACG<br>AAATCAAGtaGGTAAGCGTTTCCCGAAATCG | BP |
| FBgn0032518 | GTATTTTTTAGGTTTTTTCGTCTGCCCGTGGCAGCCACACTAATTTGGCT<br>CAGCTTTTCGTTTCCACTTCCGTTTTCTTTTCTTTTCGTGTTTCTACGCCA<br>GCAAGtaGAAGTACGTTAtaGAAAAGCGTT | NP |
| FBgn0033081 | GACACGAAATCGAGGGATGAAATTGCATGTGATGCAGCCCCTTGAGCAA<br>ACAGTGTTGGACAACAGCGCGCGGCATCGGCATTTTTTGGCGGCCATTA<br>CAACGAtaGGAAAACTGTGAAGCGTTGCGCACA | BP |
| FBgn0034010 | AGGTATCTGAAAGTCGAGACATAGTTAAGTCCACGCTTACAGATCGGGT<br>ATATAAAGAGGCCACTTTCAGAGCGGATTTCAGTTTAATAATTTAAAGCA<br>AAtaGAAACTACTGGTAAGCTAGCTGGGTTAT | NP |
| FBgn0034308 | CGCCATGCGCAGCACTATCCTGCGACTAAGCCAGATCTGACGGGAATAA<br>AGCTGAATCGGCAGCACTGCCGCAAGTATCCACTTTTTCACGGGCAACA<br>AATTGACAAAGAAATTGTAAGATAATTTCCTGT | BP |
| FBgn0034642 | TGGTGCCGATTATCTTATCGCCAAGTGTGGACTGCAAGTTGGGAAAACG<br>AATACATTCATCACCCCGGTCGTTGCTCACTAACTGGGTTTTCGGTGAC<br>GCTATTACGGACACGGACCGGCTCTCACCGAAA | NP |
| FBgn0035754 | AGTCTGGCAACCTCTCTGTTACGGTATTTTTACAACGTGGTATTAACAGC<br>GCTCCGGAATACTATACGGTATATTTCAGCAATCGAAGAACGGCCACATT<br>GCGGTGTGGAAAATAAACAAATTGCAATTAT | BP |
| FBgn0035906 | ACAATCGAAATATATTCGATAAATTCACTCGTCCGGCGACTGCGACCACT<br>TTATAAGGTACCGGAATCCCTCTATTTGTTGTCAGTCGATCGGAACTACT<br>TTGCCACCAACATTTCACGTCTTGGGAATTA | NP |
| FBgn0036263 | CTACGTAATATACTAACGCACTTTTAGGTATATTTTTCAAAAATAATATACT<br>GTTCTTGGTATATTGCTCAGGAACGGTCACTCTAGAGAGCCGGCGTAAA<br>CAAAGCGATACAATTTGGTTAAATTAATTA | BP |
| FBgn0037328 | ACACTTTCGAGCAACGGCGCGCTGTTTCACTTAACATATCGCGTTTTTGT<br>GGCGTCTAGAGGCAGCCACACTATTTCCTTCTTTTCGCTTTCGTTTCCGG<br>CGAAGTAAGTAAATTAAATTTCTCTGTATAT | NP |
| FBgn0060296 | GTGTGGCCCCTGTTAGCTTTCTGTTAAATTTAAATTTCTGTAAAGTGCCC<br>GCCACTGCGGTCGCTTTCACGGATCAGATTAGTCGTTGTCTGGATATTA<br>ACGAGGAAGGTAGTGATCGCGCATTAGTGTCA | NP |
| FBgn0064225 | TTGGCATTTATTATTTTTATTGTAAGGTATTTTTTAGTACATTTGTTTCTTG<br>GATCCAATAAGGCCGCACTATTTTCCTTCTTTTTGCTAGCAATTTCCGGC<br>GAGGTtaGTGTAAAATATTTCTACTCCAC | NP |
| FBgn0086519 | GACTTGAACTTGGGCGCCACCGGCAGAGGTAAGCTGAGCATCAGTATC<br>ATATAAAAAGCAGGCAGAAGTTCCAGTTCGATATCAGTTAGCCTTCTCAA<br>GTCTTTCAACAATCAACtaGAAGTTCTTCGTAA | NP |

**Table S3.** The sequences for 19 native core promoters investigated (with TSS downstream ATGs mutated).

| Mutation type | Number of synthetic promoters designed |
|---|---|
| knockout of motifs (individual or pairwise knockout of motifs, and knockout of all motifs) | 260 |
| replacing the original motif with sequences with different PWM scores or insertion of the consensus into the Ar.0 sequences | 170 |
| point mutation of motifs | 596 |
| substitution with functionally or positionally equivalent motifs from other architectures | 78 |
| shift of motif positions | 164 |
| intra-architectural mutations | 2023 |
| inter-architectural mutations | 478 |
| context swap | 30 |
| native sequences | 19 |
| native sequences with ATGs | 8 |
| **TOTAL:** | **3826** |

**Table S4**. Number of synthetic promoters designed for the different mutation types. Among the total number of synthetic promoters designed, ~3000 could be recovered and tested during the experimental procedure (**STAR*METHODS**).

| Block 1 | Sequence (5'-3') |
|---|---|
| 1.1<br><br>+ | CGCAGTGCAGTGAATCATCCGTGGTGACCCATGGCTCTCGACTTACAGAGCGGCTCTTGGTGTT<br>TCCCCGGTCGTAGATACACTACACTGAACGAAAATTTACGAGCCGATGCATTTACATTCCATTCC<br>ATTACATTCTCTTATATGGCATGTGCTCAATTGCTGTGGAGGATGTACGGACTAGAGATCGCCTC<br>TTTCAGTGGCGGCACACTTGGTTCGTGGTGTTTCAGTGCAGTGCT |
| 1.2<br><br>− | TTTCCAGTGTAGAAACGCGTTTATAGATGTTAAATATTAACCACTGATAAGTACAAGCTAATAACA<br>ACAATAATCGTAACAACGCGCTCTGGTTTTCTCCGTGTGCACTCAAGAGCGTTGCTGATTGAAGC<br>GGATGAAGCCGAAGCCGATCGGAGTTGGTCAATTTTCTAAACTCTCTCACGGTCTTCAATTGAAC<br>GGCACTTCCTCGACTTCCTCCCGTCGCCCCCGCCCTTTCACAC |
| 1.3<br><br>− | CGCTCCAAGCTAGACTCAAGAGAGATACGCACCGGAGATACGCAAACCGGTCGTTGGCTGGCT<br>AGCCGTAGCCAAATATTATGCTAATTCGACATTTTTGACAAAAATAATTCGAGAATTAATTATATCT<br>TTACAATGTGTCTGCTAGTTGACACATAGTTAGTTAATGTCTCCGCGTCAAACTCGTCTTCCGTCT<br>GTTTCGGGCATTATTATGGGATTCAATGCGAACTTTAACTGAA |
| 1.4<br><br>+ | TGTAAAAATTTTATTTTTGGAAAATCAAAAAACTAGTGATAGGGATAGTTAGGTATGTTTATTAGCA<br>GTACAAAACAGTCTTTATTTTCGCTGTGCCACCTTTTTTGGCCAAGTTTTGGAGAAAACCCCGTAC<br>GGGCATATCAGATTTTTAGCATCATCTTTTGCAGCGCTGGAGGAAGCCAGAGTTTTCACTCACCA<br>ATAAAAAAATGTAACTTAGTAACTTTAGCCAGATTTTCCGTT |
| 1.5<br><br>− | TGCTGCAAGTTGCTTGCTGGTTTGTATGTTTTAAGAGTGAAACTGACGGGAGACGCGAACGCGA<br>GGATGCAACAGAGTATTGTAATCTGCCATATTTGAGAAGGTTTGTTAGGTTTGATTTGGCTAATCA<br>ACAGAAGAGTTTTATGCTAAAATTGTAACTGCAATTGTAACACACGAGAAATAAACAAGCAAATAA<br>TACCTACTAGAGATAACGCTGCGAATTTGTTTTATTTTTGAACT |
| 1.6<br><br>− | CAGTTGGCGCGATAGCAGGACTTATCAATTGAATAACAGGACCTTATTATCGACAAAACCTTAGA<br>GCTGCCACGCAATTTAATAAAGTTATCGTTCGTAAAGTTATGTCGAATTTAGATTTAAATTGAAATT<br>GATGAGGTAATATATTTTTAAAATAAAATCCTATCACTTATTGTTGCTTAAACTAAAATTTGTTTCAA<br>GAAAGACTATTATGAGATAGATCTTCGACTAAAAATAAC |
| 1.7<br><br>+ | TCCCCTCTGCACCATCGTAAATATACGACTTTTATTTATTACTTCATTTTATTTTCATTATTACTATC<br>TTTGAATAAAGAAATTTCAGAATACCAAACAAAATGATTTTCGTTTCATTTTGAATTAAAATTTCAC<br>CAGTGGGGGAAAATAACGGTATTTGAACAATAATGGCGTTGCGAATGTAACGTGATTGGCGTAA<br>AATCGCAAATTCCGTTTTATATAGTTGAATGATTTTCAAA |
| 1.8<br><br>− | GCTGAAATTCCGATTTTAGGCCGCTGTCCAGTTTACAATTAATGAGATTAATCGTAAATAAACATT<br>TACGGAGAGCACTCTTACTTCTGACACACACGATAATTTGTGGCAGACACATGGAGAATGAAATT<br>TTTTATGAGGAAATCGAATTGCAATCTCCGGCGAACGATTTTGCTCATGCAATTGCAATTACAATT<br>GGTTTTCAATTGTATTTCCTCAAAATCAAAAGAATGTCGAAT |
| 1.9<br><br>− | ACCGCAGTTTCCAGCTGGCTTGAAATTTTGGCCGGCGACTCCTCCTCGTTGGCAGTTTTTAGAAA<br>ACGAATCGCCTGTGCAGAGGCCAAGGCTCCGGCCGTAAAAGAGGTCAGCAACTGAGCAATCGA<br>AATCATTTGAAATTTGATATTTTTAAATGATTGGAAAGCGAATTTAGATCATGCTTTGAAAGTTCGT<br>TACGATTGCTATGAATTGAGTTTAAATATTTCAAGGCTACATAA |
| 1.10<br><br>− | GTGGGCGTGGTGCCCGGCAAGGTGTTGTACTGCCAGTGCGGGGCCCCCAATTGCCGCCTTCGT<br>CTGCTCTAAGTTCTAGCTTAAGTTAGAGATCCATACAGGAAATATACTCATTAAAGAAGAATTAGA<br>ATTAAAAATTTAAACTTTAAACATTATCTGTTCCGTTAGGGTAACGGAAACATTGCATTTTTATAAG<br>CTACTGCTGTTCTGCACCGTCCGTTTCAAAGTACACAATTTTC |
| 1.11<br><br>− | TTGGGGGAACAGCCTGAAAGTAGGCTACAAAACTCTTGTCTTCAGTACTCTTTATCGTGATTCCC<br>ACGACGACTTTCTTGCTTTTACATCATAAACTCAATGTGGTAATAAATTTAAAAATAGTTATACTTT<br>TCTGTCATATTCACCAAAAGCTGGAAATTTGTATTAATTTTAATGTTGATATGAGGTCAAACGCAT<br>AAAATAAATGTATAAAAGATGTTTTGCTTACTCCGAAATCA |
| 1.12<br><br>+ | AATTCGGGCCTGCCCATTCAGACAGCCAGTGACTTGGATGATGCCGCCCACAAGGCTGTGGCA<br>GCCCTTAATTAGGGGAACGATTGAGGAGAGCATGTCTTCCAGAATGAAACGACGCTCATTAGCA<br>TTTACAACGGTTGGGCCTTTTAAGTATAAGTTTTTATCGACAATATAACCAAAATATGTTATATTCT<br>ATATAAAAACCTTTTTATTTGATTTAAGAAGTACCTTAGCCATCT |

**Table S5.** The sequences of block 1s used in this work.

"+" in column "Block 1" represents well-positioned -1 nucleosome pattern found in the genome-wide MNase digestion of chromatin; "**−**" represents not well-positioned -1 nucleosome pattern.

| Block 7 | Sequence (5’-3’) |
|---|---|
| 7.1<br>**–** | AGCATAGGAGCCGCACCAGGATTCGCCCGTGTACGCCAACTACGAAGATTAGCGCAACTCTG<br>GGCCGGCCAGCTCCACGGCGTACTAGGTCAACTAGGGCGCCGCCGGCTAGGCACCCGAGCC<br>GGCACTTCGAGTGCCCGGCACCACGCAGCAGTATCGTGGATTCAAAACTTGTAAGCGGGCGG<br>AAACGGTGTATACTATAGTAGCAGCAAGTAGTAGACCCATCCCGGCCAATATTAG |
| 7.2<br>**–** | GTGAAGTGCATTGAAAGCAGAGCGAATCAGAACGAAATTCAAAACGTATCACGCATACGCCCG<br>GTAGTACAGCCGAATATCCCCAAATAGCCGAATTAGTCGCCGAGAAGGAGCTTGCTGTGCTGC<br>CTGCTGGTGAGCAGCATCCTAGTGCTGCACCAGGCTCAGGCCAACATCGAGACAAACGTAGT<br>GATAGACCCCAGTTACTACAGTAAGTGTGCGTAGTGGATCAGGCGGCCATAG |
| 7.3<br>**+** | CGCAAATTGATCCTTTATAGATTCTAGCTGATTTGTTATTTACAATCCTTGTAGATTGCTTTCGC<br>CTGCCTGTTGGCCGTCGCCCTCGCCAACGAAGTAGCCATAGTTCTCCGTGCCGAGCAGCAAG<br>TGATAGTTGACGGCTTTGCTTACGCTGTTGAGCTGGACAACTCTGTCATAGTGCAACAGAAGG<br>GTGACCTTAACGGCGAAGAGTGGGTGGTGAAGGGAAGCCAGTCGTGGACA |
| 7.4<br>**+** | GGATTGATAGGCGTTATCAGTCGAAATTGAAAGGGTATAGGACCAGGGCAACTGCCTGTAGCC<br>CGACATCAATATCTGCCAAAGCGACTTGGCCAATCCCACCGAGCCCATTGTCACCAAGATCTA<br>GGTGCACTATCTGCGGAGTTTCGGCTTTCGCCTGGAGCCGCCCTATAAGATTGGCACCGAACT<br>CGGTCACTCGTCGCGGGAGGCGCGCGTCTTTCTTATCCGAGTGTGCCGCCA |
| 7.5<br>**+** | AAACGATTTGCCGTGATATAGTCGTTTCATTTGAGCACAAAAGAGCGCCTTCTGCTCCTGATCG<br>ACGACATTGAGTAGATTGCCAAGGAGTTGATCGAGCAGGCGCACCAGAAGATCTCCAGCACC<br>GAATTGGTGGACCTGTTGGATTTGCTGGTGGCCAAAGTAGAGGAATTTCGCAAATAGCTGGAA<br>CTGGCGGAGGAGCAGGCGAAAGTGGAGGAGGCGTAGGACCAACTGCGCGCT |
| 7.6<br>**+** | AAAAAATAATAAAATAGTTCACTTAAAATCCATAGCCACCTAGAATTTAGCTTGCGGGAAGCCT<br>GCATAGGAGTTTTGCCTCCGCCGCCAAGGCCGCAAAAGCAAAGCCCGTCAAAACCAGCACCA<br>TTTCAGCTGTCGGCGAGTCCATCGCCGCCAAGGGGTAAGCACTTTCTTTAATTTATTTACATAA<br>AAACCAAGTAAATATTCTCCTGATTGTAGCTTCTTGCGTCCCCATAAGCC |
| 7.7<br>**+** | CTAAAACACCGACTAAGGACCTCTGAATAGGACTCTACTGCCACTCGCACGCTGCCCTTTGTG<br>TACAAGTATAAAAATCATAAAGGCCAAGACTGCGAATCCCGGCTGGACATCACTTTTCCCTTTG<br>ACCAGGAAAATCTGGAGGAGTGTATCACCCAGCTGTAGGCTCCCAACCAGTAGGACCCTTAGT<br>AGCGGTACCTGGACGAGAATATCAGTGAGTATCAGTTAGTGCGGTAGGAG |
| 7.8<br>**+** | AAATTAGAAACAAAAGGATCTGAAACGCGCTTACAACATAGTCCTAGCACAGGCTATATCCGG<br>CTCTAAACAGTACCTATTTGCGGGAAATCTCTTTGGCGACATTTTCGTACTTAGGTGAGAATCC<br>TTAATTTGGCGCAATTTCCCAACTAATTGTTTTTCTATTTTGAAGAATAAAAGAACTGGACAAGG<br>GTTCCGAGGAGCCACCTGGCAAACTGAAGATCTTCCCGCAGGGCAGCG |
| 7.9<br>**+** | AACTTTATAGAGTTTTATCCAATTTGAGCGTAGCTGCCGGGCGTTGGAGTGTTCGGAACGGGA<br>GAGATAGCCATAGTGCTGGTGCCGCTGCTGCGGGAGAAAGGATTCGAGGTGCGGGCCATTTG<br>GGGCAGGACCCTGAAAGAGGCGAAAGAGACTGCGACCACGCAGATAGTACAATTCCATACGA<br>ACGTAATCGACGTAGTCCTGCTGCGGAAGGTAGTGGATCTGGTGTTCATCGTG |
| 7.10<br>**+** | ATTCGAGAGTAGCTAAATTAGGGGCCGCATAGCGCCAAGCTCACCTGTAAGTGGCTCCAGCAA<br>GTGTCTCGTCGCAGGTGCACAGGTGCTAGGCAGCAGGACTCCGCCGACGGCTGACTAAGAG<br>CAGCTTATCCGCCCCTTGGGAAGAAAAAATCGATTTTGTGATAACGGGCTTACGCTTACGCGT<br>TTTTGCACTGCTAGTCGGAACCAATTGCCAGTAGACTCCACACCTAGATAGTG |
| 7.11<br>**+** | CGATTGCACACGTTGCACTTTTGTTGGCATTGGACGAGGTCGAGTGAAAGCACAGCGCAAGC<br>CCCGAGAATCCCGATTCGTTTGCTTAGTTCAGGGTCAGGCCGCTCCTCCCCAGGATACGAAG<br>CTCCCTGCAAACGACCTTGAAACTCCAAGCAGATACATCGCAATCCGAATCCGAATCGTTCCG<br>TCGGAACCTACGACTCCTACACAACAGTTGTCGTAGTCGCCCTGTGAGCAAGT |
| 7.12<br>**+** | TAGAATTTCCCACTCAAAGGTAGCAAAAACCATCCCACTTCGCACGCGAGGATTCCCAGGAGC<br>ACCTGCCCACCAGCCACACGCAAAACAGTCACACAGTGAACTAGAAGAGACGCACACTATCC<br>GGAAGTTGTGGCGTTGGCGTCTTCGTGTTCGCCTTTGCCTTCATCGTGATTGCGTTTGCAACG<br>CCCAGTTGGTTGGTCAGTGATTACCGCATCACGGGCGCCAAGCTGGATCGCC |

**Table S6.** The sequences of block 7s used in the thesis**.**

“+” in column “Block 7” represents well-positioned +1 nucleosome pattern found in the genome-wide MNase digestion of chromatin. “**–**” represents not well-positioned +1 nucleosome pattern.

| Block | Coefficient | Standard Error | t-statistic | *p*-value | Significance |
|---|---|---|---|---|---|
| (Intercept) | -2.65885 | 1.36776 | -1.944 | 0.053117 | . |
| Block3_FBgn0031980N | -1.57372 | 0.61621 | -2.554 | 0.011297 | * |
| Block3_FBgn0034010 | -2.73639 | 0.62579 | -4.373 | $1.86×10^{-5}$ | *** |
| Block3_FBgn0064225 | 0.15807 | 0.61497 | 0.257 | 0.797373 | |
| Block3_FBgn0086519 | -2.20657 | 0.61994 | -3.559 | 0.000451 | *** |
| Block4_FBgn0004878 | 0.09943 | 0.26785 | 0.371 | 0.710835 | |
| Block4_FBgn0031980N | 1.75593 | 0.27639 | 6.353 | $1.11×10^{-9}$ | *** |
| Block4_FBgn0034010 | 1.88443 | 0.27366 | 6.886 | $5.37×10\text{-}^{11}$ | *** |
| Block4_FBgn0036263 | 0.19417 | 0.26369 | 0.736 | 0.462259 | |
| Block4_FBgn0064225 | NA | NA | NA | NA | |
| Block5_FBgn0031980N | 0.35498 | 1.4742 | 0.241 | 0.809927 | |
| Block5_FBgn0034010 | 0.38067 | 1.47562 | 0.258 | 0.796657 | |
| Block5_FBgn0036263 | 2.07272 | 1.47939 | 1.401 | 0.162536 | |
| Block5_FBgn0064225 | -0.35306 | 1.48039 | -0.238 | 0.81171 | |
| Block6_FBgn0004878 | 1.97382 | 0.37046 | 5.328 | $2.35×10^{-7}$ | *** |
| Block6_FBgn0014865 | NA | NA | NA | NA | |
| Block6_FBgn0031980N | -0.05506 | 0.25839 | -0.213 | 0.831442 | |
| Block6_FBgn0034010 | -0.70079 | 0.26107 | -2.684 | 0.007793 | ** |
| Block6_FBgn0036263 | -0.07401 | 0.29249 | -0.253 | 0.800463 | |
| Block6_FBgn0064225 | -0.14806 | 0.26584 | -0.557 | 0.578106 | |

**Table S7.** Coefficients of the linear regression model for the inter-architectural block-wise combinatorial mutations.

Significance level codes: $p \leq 0.1$; *$p \leq 0.05$; **$p \leq 0.01$; ***$p \leq 0.001$. NA: not defined because of singularities.

### Supplemental Texts

**Supplemental Text 1.** Various potential nucleosomal contexts affect expression

We checked the influence of different nucleosomal contexts on the expression level of five native core promoters (Mtk, RpL23, Mec2, CG17712, and geminin) selected from the different architectures, and such that their activities covered the entire dynamic range of our measurements. We created constructs containing these promoters surrounded by all free combinations of the different available *blocks 1.X* and *7.X* (**Figure S3C; Tables S5** and **S6**). In total, we tested 127 promoter variants recovered out of 136 possible combinations of different *block 1*s and *7*s. As expected, we observed lower activities on average for the five native promoters compared to the constructs containing combinations *B1.11* + *B7.11* (an average signal reduction > 2.5-fold), the fixed pair of nucleosomal sequences selected for the most of our constructs. A more detailed promoter-wise analysis (**Figure S3D – S3E**) revealed smaller variations of the expression levels for *B1.X* + *B7.11* samples compared to *B1.11* + *B7.X*. This indicates that *block 7*s might have relatively stronger influences than *block 1*s on the expression level. The sequence downstream the TSS (*B7.X*) forming a prominent +1 nucleosome may set a transcriptional obstacle. It may also influence post-transcriptional events as it constitutes the main component of the 5' UTR region. We computed the GC content of each *block 1*s and *7*s, speculating that as the GC content usually correlates well with nucleosome occupancy, it might correlate with our expression data. We however could not find any clear relationship between GC content of the different *block 1*s and *7*s and the expression levels (**Figure S3C**). Furthermore, to test if the presence or absence of *block 7*s has a strong influence on the expression levels, we also checked core promoter sequences with *block 1*s variants only, and no *block 7* (**Figure S3F**). In these constructs, the length of the 5' UTR was thus reduced from 333 nt to 89 nt. We observed that the expression levels of promoters with or without *block 7.11* sequence, were in the same order of magnitude, whether induced by ecdysone or not (**Figure S3F**; all constructs contained *block 1*.11 kept constant, *PCC* $r = 0.96$, $p = 1.2\times10^{-5}$). Hence, *block 7* presence and its content have an influence on expression level, but the effect is limited compared to core promoter motifs.

After having evaluated the overall effect of different nucleosomal sequences, we next explored potential promoter specificity. Indeed, the two tested constitutive promoters RpL23 and CG17712 exhibited stronger expression variations when altering *block 7s* (the median SD within the same *block 1* is 1.23 compared to 0.66 for developmental promoters; Wilcoxon rank-sum test $p = 3.1\times10^{-4}$; **Figure S1D**, lower panel). In contrast, both the constitutive and developmental promoters showed similar and milder expression fluctuations upon *block 1* variation (the median SDs within the same *block 7* for constitutive and developmental promoters are 0.64 and 0.54, respectively; Wilcoxon rank-sum test $p = 0.3$, not significant; **Figure S1E**, lower panel). Since previous genome-wide studies showed constitutive promoters tend to have a preferred canonical nucleosome pattern with a strongly positioned +1 nucleosome ((Mavrich et al., 2008; Rach et al., 2011), *block 7s* which were designed in our experiments to act as different potential +1 nucleosomes could have more prominent influences on constitutive promoter activities.

**Supplemental Text 2.** Linear regression analysis for inter-architectural block-wise combinatorial mutations.

We built constructs with random selections of the individual *blocks 3, 4, 5,* or *6*, respectively, for 5 different promoter architectures (**Figure 3D** and **STAR*METHODS**). Similarly to the analysis of the intra-mutations, a linear regression model was learned directly from the expression measurements obtained from these block-wise combinatorial mutations (**Figure 6B**; detailed mutation design in **STAR*METHODS**). The predicted values also showed a high correlation with the measured expressions (*PCC* $r = 0.81$, $p < 2.2\times10^{-16}$), supporting the additivity for sequence features even among various promoter architectures. The learned coefficients revealed the significance of the block features (**Table S7**), although some coefficients were not significant, probably due to too sparse data (not all inter-architectural mutated promoter constructs were recovered during the cloning procedure; **STAR*METHODS**). Surprisingly, *Block 5 sequence*s generally had a weak impact on the predictions (average $p > 0.6$; **Table S7**). As the tested *block 5* sequences always contain functionally similar motifs essential for transcription initiation (such as INR, INR2, CA-INR, Ohler7 and R INR), our results indicate that binding to these motifs is retained, even by exchanging *block 5*. However, one cannot exclude that this block feature is closely correlated to other blocks due to lack of data (only ~50% of the sequences were recovered for the inter-architectural mutations during the experimental procedure), leading to a non-significant impact on prediction. Ignoring *block 5*, the block variants with the strongest contributions to expression correlated with the influence on expressions of specific motifs inside these blocks. For instance, *block 4* in CG8157, *block 4* in RpL36AN and *block 6* in Cas were the most significant features (p < 3×10-7; **Table S7**) found in the model, in which TATA-Box, DRE and MTEDPE motifs locate, respectively. They all increased the expression levels when replacing other blocks (average coefficient > 1.85). *Block 3* in RpL36AN, which contains DRE, gave a negative contribution (coefficient = -1.57, $p$ = 0.011). This indicates a possible positional preference for DRE to be located in *block 4*. The background sequences of *block 3* in CG8157 and Cpr47Eg (with a non-functional CGpal) provided significantly negative effects (average coefficient < -2.4). *Block 6* in CG8157 with a TTGTTrev played a slightly negative role as well, which is again consistent with the repressive function of this motif (coefficient = -0.7, $p$ = 0.008).
